# Rat microbial biogeography and age-dependent lactic acid bacteria in healthy lungs

**DOI:** 10.1101/2023.05.19.541527

**Authors:** Lan Zhao, Christine M. Cunningham, Adam M. Andruska, Katharina Schimmel, Md Khadem Ali, Dongeon Kim, Shenbiao Gu, Jason L. Chang, Edda Spiekerkoetter, Mark R. Nicolls

**Author notes:** To whom correspondence should be addressed Lan Zhao, PhD Division of Pulmonary, Allergy & Critical Care Medicine School of Medicine Stanford University 1701 Page Mill Road, Palo Alto United States.

## Abstract

The laboratory rat emerges as a useful tool for studying the interaction between the host and its microbiome. To advance principles relevant to the human microbiome, we systematically investigated and defined a multi-tissue full lifespan microbial biogeography for healthy Fischer 344 rats. Microbial community profiling data was extracted and integrated with host transcriptomic data from the Sequencing Quality Control (SEQC) consortium. Unsupervised machine learning, Spearman’s correlation, taxonomic diversity, and abundance analyses were performed to determine and characterize the rat microbial biogeography and the identification of four inter-tissue microbial heterogeneity patterns (P1-P4). The 11 body habitats harbor a greater diversity of microbes than previously suspected. Lactic acid bacteria (LAB) abundances progressively declined in lungs from breastfeed newborn to adolescence/adult and was below detectable levels in elderly rats. LAB’s presence and levels in lungs were further evaluated by PCR in the two validation datasets. The lung, testes, thymus, kidney, adrenal, and muscle niches were found to have age-dependent alterations in microbial abundance. P1 is dominated by lung samples. P2 contains the largest sample size and is enriched for environmental species. Liver and muscle samples were mostly classified into P3. Archaea species were exclusively enriched in P4. The 357 pattern-specific microbial signatures were positively correlated with host genes in cell migration and proliferation (P1), DNA damage repair and synaptic transmissions (P2), as well as DNA transcription and cell cycle in P3. Our study established a link between metabolic properties of LAB with lung microbiota maturation and development. Breastfeeding and environmental exposure influence microbiome composition and host health and longevity. The inferred rat microbial biogeography and pattern-specific microbial signatures would be useful for microbiome therapeutic approaches to human health and good quality of life.

**Graphical Abstract:** 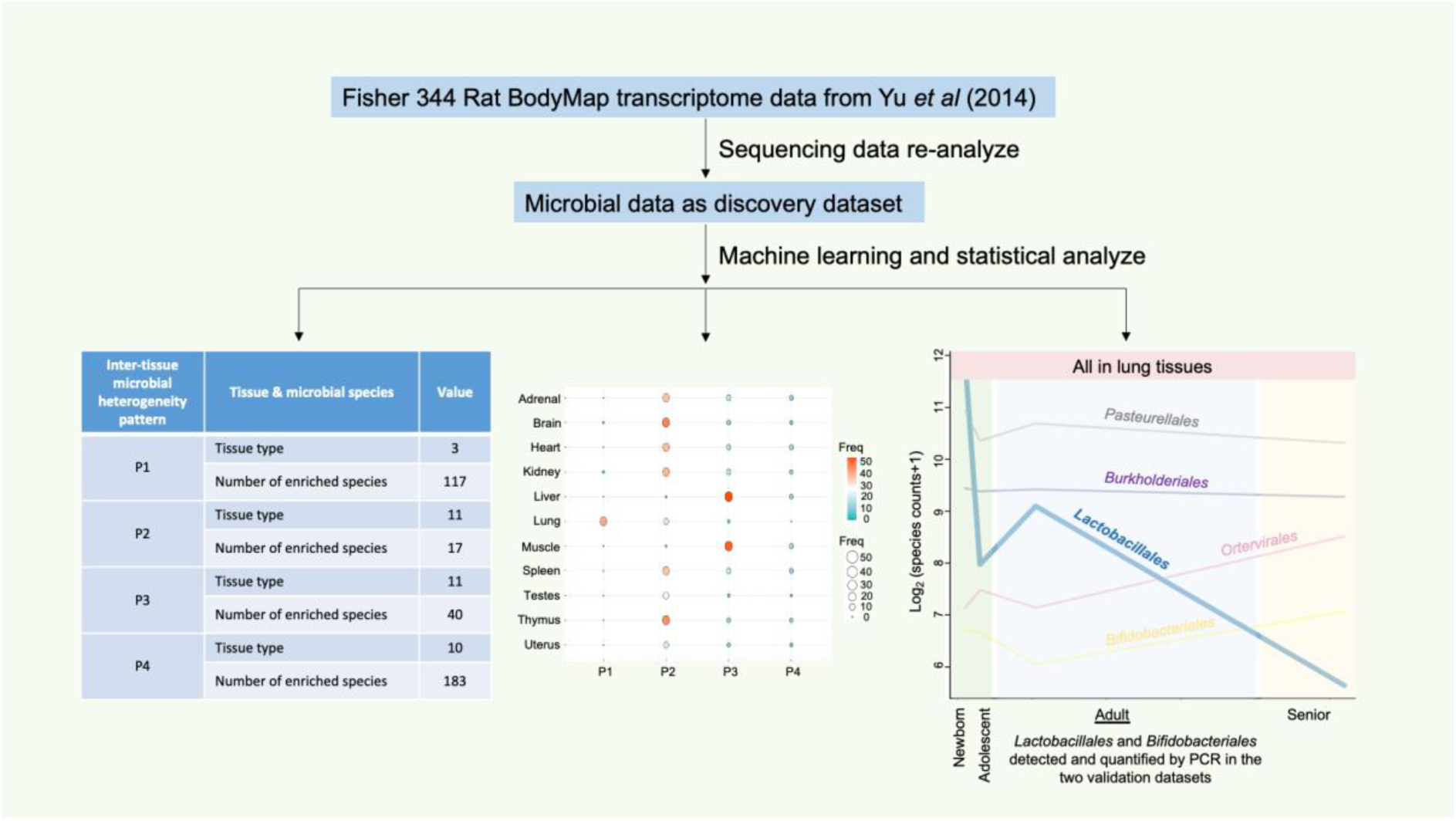

## 1. Background

The laboratory rat has been widely used and well examined as a model in a variety of biomedical fields, from cardiovascular diseases to cancer [1]. Recent evidence from the Human Microbiome Project (HMP) [2] and The Cancer Genome Atlas (TCGA) pan-cancer microbiome projects [3,4] suggest that different body sites and disease status feature distinct microbial communities which play essential roles in human physiology, health and disease. In this paper, we examine the spatial and longitudinal structures of the microbial community in various body compartments and across different life-history stages, in order to characterize the microbiota landscape of Fischer 344 (F344) rat with a view to help advance human microbiome research.

The F344 is an inbred laboratory strain of rats that is frequently used in aging, cancer, and toxicity studies [5]. Throughout the natural lifespan of F344, 2 to 104 weeks would be equivalent to 1-3 months to 70-80 years in humans [6]. In F344 male and female rats, weaning normally occurs at the age of week 3 and sexual maturity by week 7 of age. Thus, our microbial discovery cohort which was based on the SEQC data [7], consecutively constitutes four major life-history stages of rats: newborns (2-week-old), adolescents (6-week-old), adults (21-week-old), and seniors (104-week-old). The NIH’s Biology of Aging Program (BAP) used multiple mammal and non-mammalian model systems, including F344 rats, to investigate genetics and other aging-related degenerative changes, but how microbes might be involved in these processes in F344 are unknown.

In mammals, microbial colonization starts in utero and extends throughout the lifespan, particularly in newborn infants who experience rapid microbial community changes. Human placenta harbors a unique low-abundance microbiome composed of commensal bacteria such as *E. coli*, *Prevotella tannerae*, and *Neisseria* spp. [8]. Maternal-fetal transmission of microbes has taken place long before birth [9]. Newborns are then exposed to microbes from birth, during breastfeeding, and through interactions with their surrounding environments that colonize the newborn’s skin [10], oral cavity [11], gut [12], respiratory tract [13], and other mucosal surfaces [14]. For example, the neonatal skin microbiome is dynamic, site-specific, and varies from individual to individual [10]. Neonatal oral microbiota is dominated by *Streptococcus* within the *Firmicutes* phylum [11]. *Bifidobacterium* and *Lactobacillus* spp. are among the first colonizers of the gastrointestinal (GI) tract, representing the pioneer microbial communities in the newborn’s gut [12]. The replacement of breast milk or infant formula with solid foods greatly changes the infant’s microbial composition to resemble an adult-like gut microbiota [15]. Each microbial niche in the body tends to develop and mature both independently and cooperatively beginning in the first stages of life [13]. Human microbiomes, particularly gut microbiomes, remain stable once established during adulthood [16]. Later in life, elderly subjects are characterized by decreased microbial diversity and shifts in community structure [17]. For laboratory neonatal rodent models, which are more exposed to fecal and environmental contaminants than humans [18], with limited studies to show that *Enterobacteriaceae* and *Lactobacillus* are among the most dominant bacteria in newborn mice and rats [19,20]. Similar to studies carried out in humans, adult rodents are typically sampled where the microbial community is more stable.

Translocation of indigenous viable microbes, such as *Enterobacteriaceae*, *Lactobacillus*, and *Staphylococcus* [19], from the GI tract to other distant organs via systemic circulation is common in humans and rats [19,21]. Furthermore, microbial bidirectional interactions across multiple organs, including (but not limited to) the gut-brain, gut-liver, and gut-lung-axes, are important for colonization and cross-niche microbial ecosystems [22–24]. Although the gut microbiome is the largest reservoir of microorganisms in mammals, the current study expands on other organ systems to provide more in-depth analyses on currently unknown microbial biogeography in different niches of the rat body. More specifically, we focused our attention on this RNA-Seq-derived rat microbial data from 11 organs including brain, lung, heart, liver, muscle, spleen, thymus, kidney, adrenal gland, uterus (females), and testes (males) [7]. We take advantage of this longitudinal data to investigate the tissue-level microbial diversity and community changes during the full lifespan of rats, and aim to identify age-dependent species for future experimental validations. What’s more, inter-tissue microbial heterogeneity and microbe-host gene interactions were well examined and assessed. Our aims were to: (1) identify core and uncommon microbial species in healthy rat, (2) determine and compare microbial profiles of different anatomical sites over time, (3) define age-dependent microbial species, (4) investigate inter-tissue microbial heterogeneity, and (5) build microbe-host gene interaction networks in F344 rats. In addition to characterizing the distribution of microbes and their phylogenetic diversities spatially and longitudinally, we also identified niche-specific microbial signatures and patterns for establishing future links between microbiome and human health.

## 2. Materials and methods

### 2.1 Human lung tissue collection, DNA extraction, and PCR

Our study included three male participants who were donors of lung at the Stanford Hospital (CA, USA) in the year of 2022 were recruited, and all participants signed a written informed consent. The study was performed in accordance with the Declaration of Helsinki. The Ethics Committees of Stanford Health Care approved the study protocol. Demographic data including age, ethnicity, and body mass index (BMI) were summarized in Table 1. Right lung tissues were sampled and embedded in Formalin-Fixed Paraffin-Embedded (FFPE) blocks. Genomic DNA was isolated from archived FFPE using the DNeasy Blood & Tissue Kit (QIAGEN) following the manufacturer’s protocols. DNA concentration and purity were measured by the Nanodrop ND-1000 spectrophotometer.

**Table 1.**
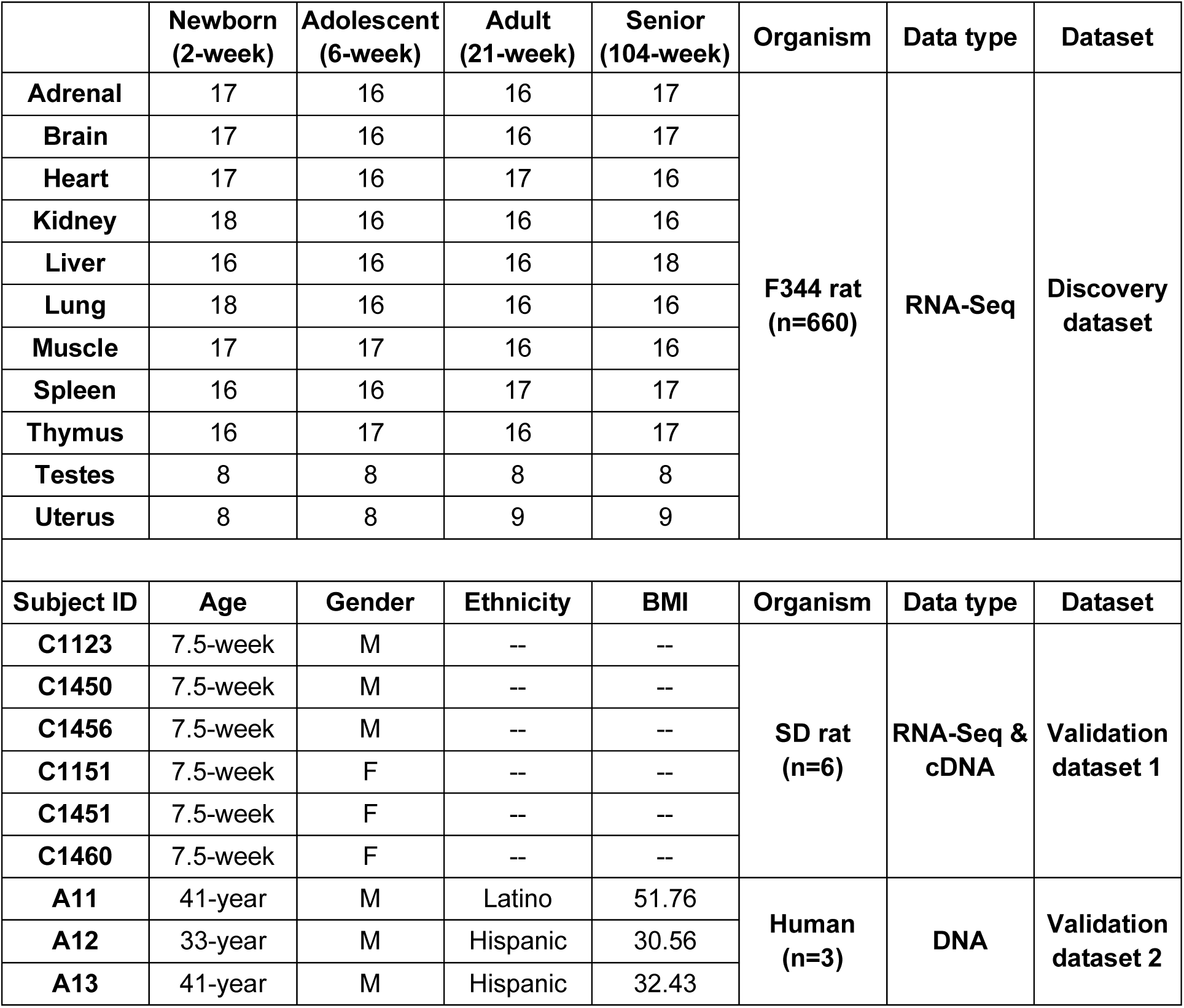
Summary of the datasets and sample sizes of the study populations.

The DNA was then subjected to conventional PCR using JumpStart REDTaq ReadyMix PCR Reaction Mix (SIGMA) and genus-specific primers for bacteria detection in human lung tissues. The primers were taken from Delroisse *et al* [25] which allow for the detection of a wide range of *Lactobacillus* spp. (15 species) and *Bifidobacterium* spp. (30 species) through PCR amplification. Only forward and reverse primers were used and synthesized by Elim Biopharmaceuticals (CA, USA). PCR reactions without primer served as negative controls. The PCR amplification protocol was described in [25], and PCR products were evaluated by 2% agarose gel electrophoresis.

### 2.2 Rat lung tissue collection, real-time PCR, and RNA sequencing

Six Sprague Dawley (SD) rats, including three males and three females (Table 1), from Charles River were housed at the VA Palo Alto Health Care System facility under a 12-hour light/dark cycle in standard conditions. Rats were fed with a normal grain-based chow diet and water ad libitum. The rats were sacrificed at 7.5 weeks to collect lung tissues. Tissues were snap frozen in liquid nitrogen immediately after excision and stored at –80°C. Total RNA was extracted using the RNeasy Plus Mini Kit (QIAGEN) following the manufacturer’s instructions. The study had ethical approval from the Ethics Committee of IACUC.

RNA qualities were measured by Agilent 2100 Bioanalyzer. The RNA integrity numbers (RINs) of the six RNA samples were all above 8. Reverse transcription was performed using the High Capacity cDNA Reverse Transcription Kit with RNAse Inhibitor and random primers (Applied Biosystems) under manufacturer’s recommended conditions. Single-stranded cDNAs were then subjected to SYBR Green real-time PCR (RT-PCR) using primers described in section 2.1 for the detection and quantification of *Lactobacillus* and *Bifidobacterium* spp. in rat lungs.

The rat *Actb* gene was used as the house-keeping reference, and reactions without cDNA template served as negative controls. RT-PCR reactions (all samples are run in triplicate) were carried out on the QuantStudio™ 7 Flex system (Applied Biosystems), and results were calculated by the delta-delta Ct (ΔΔCt) method.

PolyA enriched, 150 bp paired-end mRNA-Seq libraries were prepared, and high-throughput sequencing was performed on Illumina NovaSeq platform by Novogene (CA, USA). Sequencing data has been deposited to the European Nucleotide Archive (ENA) under accession number PRJEB57257.

### 2.3 RNA-Seq data acquisition and processing

Raw RNA-Seq data were downloaded from the ENA database under accession number PRJNA238328. The longitudinal data was generated by the SEQC consortium from 11 organs of both sexes of F344 rats [7]. More specifically, the 11 organs were brain, lung, heart, liver, muscle, spleen, thymus, kidney, adrenal gland, uterus, and testes; four developmental stages from newborns (2-week-old), adolescents (6-week-old), adults (21-week-old), to seniors (104-week-old) were included in our analyses. Rats were fed with a cereal-based NIH-31 diet (ad libitum). Housing, necropsy, organ collection, and sequencing can be found in the original article [7]. A total of 660 samples were collected and sequenced from 32 healthy rats. There were four females and four males in each organ and age group with two to four technical replicates. The sample sizes and grouping categories were summarized in Table 1.

Sequences including the PRJNA238328 and our newly generated RNA-Seq data were mapped to the Rnor 6.0 reference genome (Ensembl release 104) using STAR (v2.7.9a). Uniquely mapped reads were counted to each gene and converted to transcripts per million (TPM) for subsequent correlation analysis. The unmapped reads were subjected to re-mapping against a Kraken database, which was built from complete Bacterial, Viral, and Archaeal reference genomes from RefSeq [26].

Kraken (v2.0.8-beta) species-level taxonomic assignments for all samples were combined, and the phyloseq R package (v1.36.0) [27] was used for the phylogenetic and diversity analyses. Core microbial species were identified using a prevalence threshold of 1.0 (presence in all samples). Rare species that were not present in at least one read count in 10% of the samples were removed from further consideration. Median sequencing depth normalization was used for correcting sequencing depth differences between samples. Microbial count data were log2-transformed before statistical analyses.

### 2.4 Statistical analyses of microbial community data

Differences in microbial communities within groups (alpha diversity) and between groups (beta diversity) were evaluated and compared at phylum, order, and species levels, respectively. Alpha diversity within a sample was measured using the Shannon index, and the diversity differences between groups were assessed using the pairwise Wilcoxon rank sum tests. Benjamini-Hochberg (BH) correction was used to adjust for multiple comparisons. An adjusted p-value < 0.05 was considered statistically significant. Principal coordinates analysis (PCoA) and permutational multivariate analysis of variance (PERMANOVA) were performed using the Bray-Curtis distance with 999 permutations to evaluate differences in beta diversity of microbial communities. The analysis was carried out using adonis in the vegan package [28]. Diversity differences between groups were assessed by the pairwise multilevel comparison using adonis with the Bonferroni correction, and p-values of less than 0.05 were considered significant. The PCoA was visualized interactively in three-dimensions with the rgl package [29].

Kruskal’s test with FDR adjustment for multiple comparisons was used to evaluate the correlation between microbial species and grouping variables, such as age groups and tissue groups. Statistical significance was set as p-value < 0.05.

### 2.5 Identification of rat inter-tissue microbial heterogeneity patterns

Microbial species that were present in at least one read count in 10% of the total samples (n=660; discovery dataset) were used for the subsequent unsupervised clustering. Consensus non-negative matrix factorization (cNMF, v1.3), which enables the identification of biologically meaningful patterns in high-dimensional datasets [30], was applied with Kullback-Leibler divergence (KL-divergence) to identify the inter-tissue microbial heterogeneity patterns and signatures in the study.

cNMF clustering was run on values of K from 2 to 10, and the value of K that maximizes stability was selected as the optimal cluster number. Outliers were filtered out based on K-nearest neighbor (KNN) imputation, and ‘Max’ method [31] was used for microbial signature selection for each pattern.

### 2.6 Construction of microbe-host gene interaction networks

A microbe-host gene interaction network consists of a collection of microbial species and their correlated host genes. Spearman’s correlation analyses were performed to examine the correlations between microbial abundances of each species in each pattern with host transcriptomes. FDR adjusted p-values of < 0.05 were considered statistically significant. The union of all selected significantly correlated genes in each pattern were used to construct a microbe-host gene heatmap, respectively, for each pattern. Hierarchical clustering of microbes and genes was done using the complete method with Euclidean distances (default settings) in the pheatmap package in R [32]. Absolute Spearman’s correlation coefficients no less than 0.3 were considered further for the following functional enrichment analysis.

### 2.7 Functional enrichment analysis

BiomaRt [33] was used to perform gene nomenclature and rat-human ortholog conversions. KEGG and Gene ontology (GO) enrichment analysis of the selected genes (in human gene symbols) significantly correlated with microbial species were analyzed by using Enrichr [34], and the resulting adjusted p-values of smaller than 0.05 were considered significant.

## 3. Results

### 3.1 Core microbial species in healthy rats

Core microbial species can be defined as a group of species that are present in all investigated samples, which might be biologically associated with particular habitats [35]. In the F344 rat dataset (n=660; Table 1) [7], we identified a total of 17 core bacterial species in 4 major phyla (including *Firmicutes*, *Proteobacteria*, *Actinobacteria*, and *Bacteroidetes*) and 2 *murine sarcoma viruses* (Figure 1a-b). *Cutibacterium acnes*, *Escherichia coli*, *Pasteurella multocida*, *Bifidobacterium adolescentis*, and 3 *Staphylococcus* spp. (*S. aureus*, *S. cohnii*, and *S. haemolyticus*) are part of the commensal flora of human and animal microbiomes [36]. The *Harvey* and *Kirsten murine sarcoma virus* (*Ha-MuSV* and *Ki-MuSV*) are two endogenous leukemia viral sequences detected in both tumor and normal tissues in rats [37]. 10 of the 19 core species are environmental species, which including 6 soil bacterium (*Bacillus cereus, Sinorhizobium meliloti*, *Ralstonia solanacearum*, *Clostridium botulinum*, *Streptomyces lividans*, and *Spirosoma pollinicola*) and 4 water sediment-associated microorganisms (*Exiguobacterium* sp. N4-1P, Halomonas sp. JS92-SW72, *Hydrogenophaga* sp. NH-16, and *Alcanivorax* sp. N3-2A).

**Figure 1.**
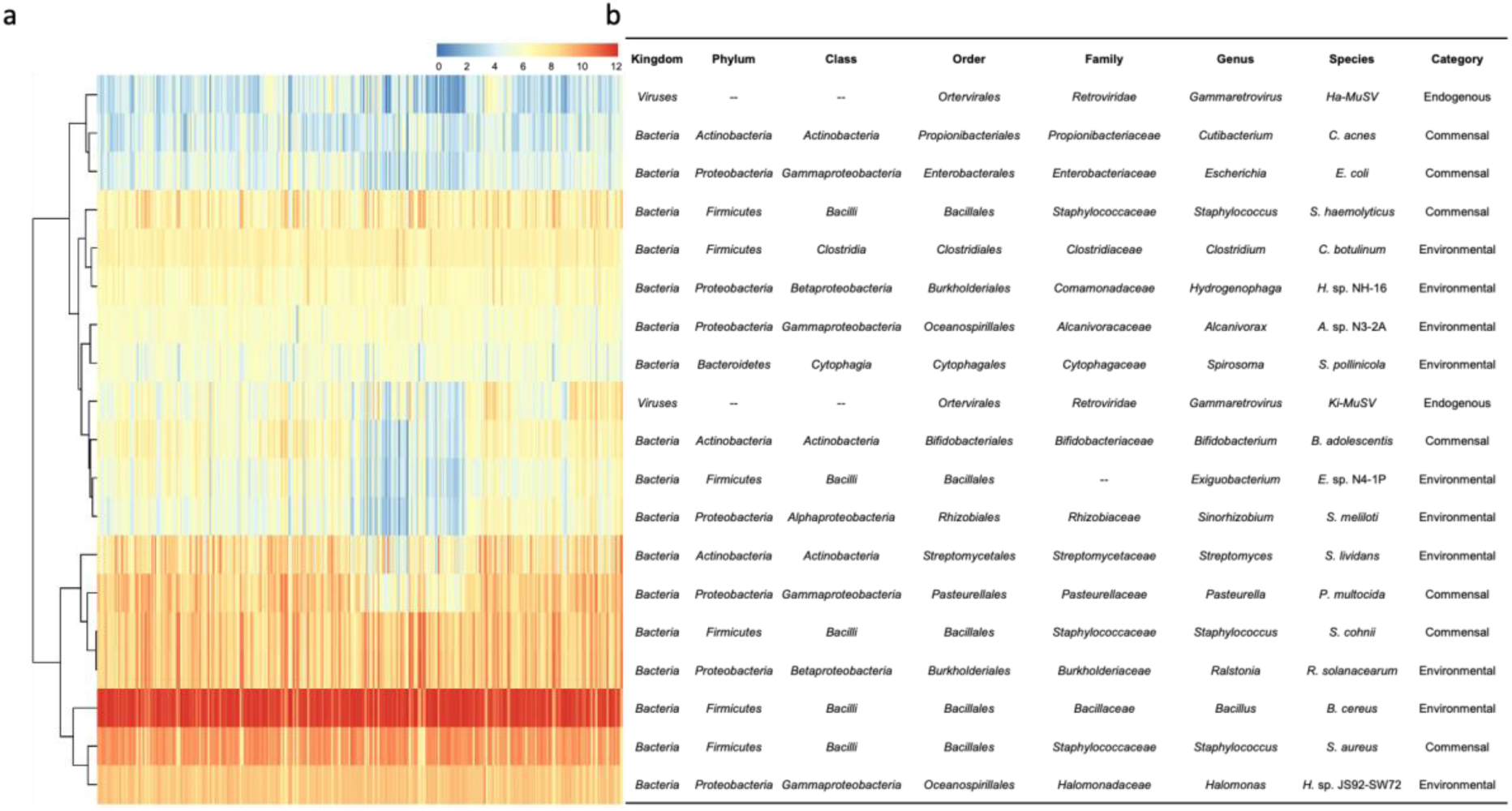
19 core species in the developing microbiota of healthy rats. (a). Heatmap of the abundance of the 19 core species in the 660 samples. The columns represent the samples, and rows are the species. Red indicates higher abundances and blue represents lower abundances. The samples and species were hierarchically clustered (complete linkage, Euclidean distance). (b). Taxonomy ranks information of the 19 core species.

Hierarchical clustering of the 19 core species identified two major groups: one with low and one with high abundance (Figure 1a). *Ha-MuSV*, *C. acnes*, and *E. coli* were among the lowest abundances across all samples, compared to the other core species. *Ki-MuSV*, *B. adolescentis*, and two environmental species–*Exiguobacterium* sp. N4-1P and *S. meliloti* showed relatively lower abundances mostly in a subset of liver and muscle samples (Table S1). The most abundant core species across all rat tissues was *B. cereus (*Figure 1a). Previous studies have shown beneficial effects of *B. cereus* on the health of animals [38], and one of the *B. cereus* var. Toyoi has been authorized to be used as feed supplements in piglet farming [39]. The pathogenic potential of *B. cereus* is well recognized in humans recently [40], thus safety practices should always be adopted when working with (laboratory) animals.

### 3.2 Microbial composition and diversity of the four developmental stages in rats

We identified a total of 2829 microbial species in the F344 rat dataset (n=660) spanning 45 unique and unclassified phyla at a 10% prevalence threshold (Table S2). *Firmicutes* and *Proteobacteria* were the two dominant phyla, altogether ranging from 86.8% in seniors to 89.4% in adolescent rats (Figure 2a; Table S3). The other phyla include *Actinobacteria, Euryarchaeota,* and *Bacteroidetes* (Figure 2a; Table S3). *Bacillales* from the phylum *Firmicutes* was the most abundant order, followed by *Burkholderiales*, *Pasteurellales* and *Oceanospirillales* from *Proteobacteria* (Figure 2b; Table S3). Although microbial community compositions at both phylum and order levels showed no significant difference among the four developmental stages, 11 bacteria were identified to be associated with the stage at the species level as determined by Kruskal’s test adjusted for multiple comparisons (Table 2; FDR adjusted p<0.05).

**Figure 2.**
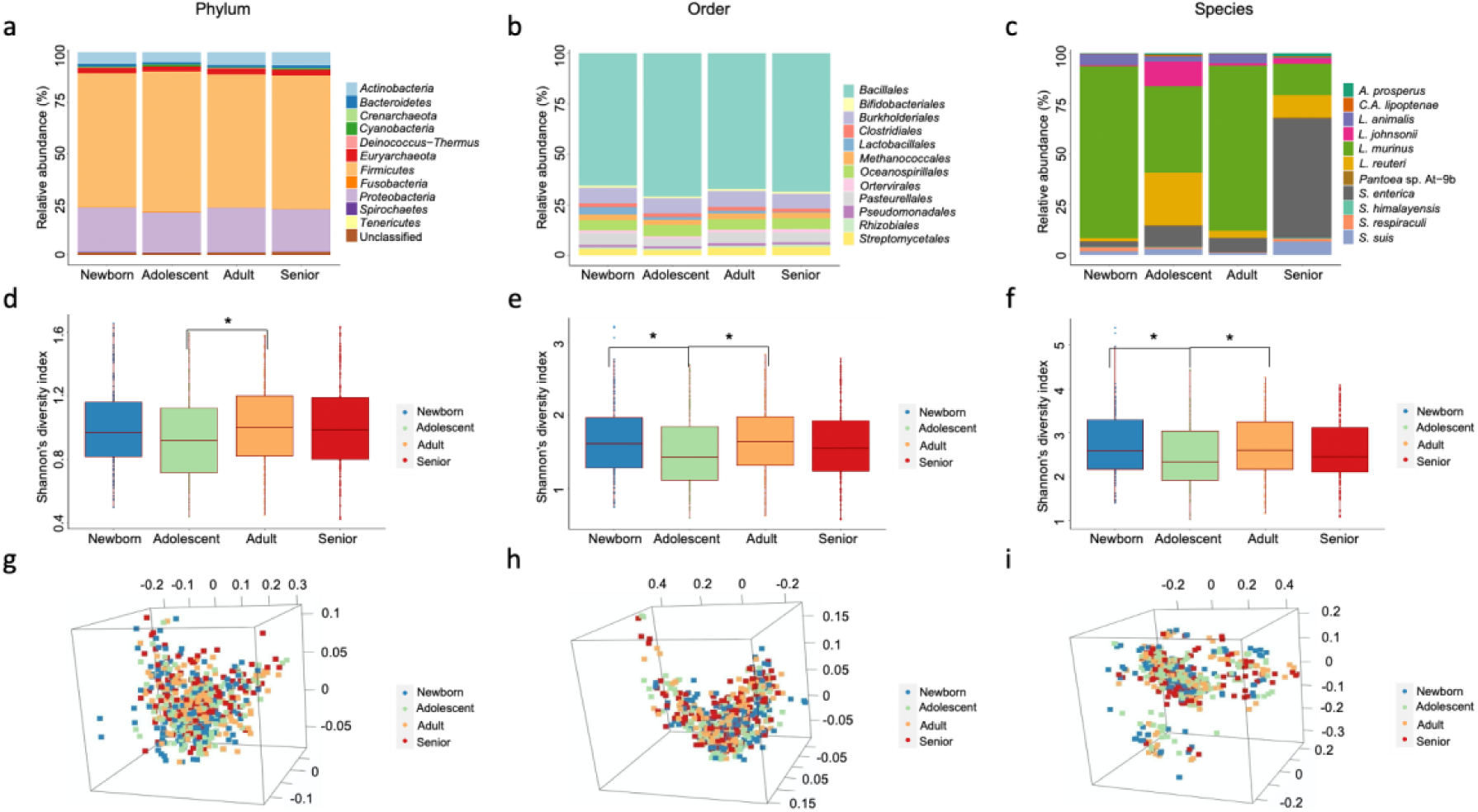
Microbial composition and diversity comparisons of the four developmental stages in rats at different taxonomy ranks. Taxa composition bar plots illustrate the microbial relative abundance (%; in y-axis) of the top 12 most abundant taxa of the four developmental stages at the phylum (a) and the order level (b). Relative abundance (%; in y-axis) of the 11 stage-related bacteria of the four developmental stages at the species level (c). Shannon’s diversity index (in y-axis) of the four stages at the phylum (d), the order (e), and the species level (f). Pairwise Wilcoxon rank sum tests with BH correction were used to test for diversity differences between stages. Only statistically significant comparisons (adjusted p < 0.05) were marked with a single star (*), and adjusted p-values < 0.01 were marked with two stars (**). 3D-view PCoA of Bray-Curtis dissimilarity between samples colored in four different colors according to each developmental stage at the phylum (g), the order (h), and the species level (i).

**Table 2.** Taxonomy information of the 11 age-dependent species.

| Adjusted p-value | Kingdom | Phylum | Class | Order | Family | Genus | Species |
| --- | --- | --- | --- | --- | --- | --- | --- |
| 0.043 | Bacteria | *Proteobacteria* | *Gammaproteobacteria* | *Chromatiales* | *Ectothiorhodospiraceae* | *Acidihalobacter* | *A. prosperus* |
| 0.022 | Bacteria | *Proteobacteria* | *Gammaproteobacteria* | *Enterobacterales* | *Enterobacteriaceae* | *Salmonella* | *S. enterica* |
| 0.030 | Bacteria | *Proteobacteria* | *Gammaproteobacteria* | *Enterobacterales* | *Erwinaceae* | *Pantoea* | *P. sp. At-9b* |
| 0.036 | Bacteria | *Proteobacteria* | *Gammaproteobacteria* | *Enterobacterales* | *Morganellaceae* | *Arsenophonus* | *C. A. lipoptenae* |
| 0.001 | Bacteria | *Firmicutes* | *Bacilli* | *Lactobacillales* | *Lactobacillaceae* | *Lactobacillus* | *L. animalis* |
| 0.014 | Bacteria | *Firmicutes* | *Bacilli* | *Lactobacillales* | *Streptococcaceae* | *Streptococcus* | *S. himalayensis* |
| 0.014 | Bacteria | *Firmicutes* | *Bacilli* | *Lactobacillales* | *Lactobacillaceae* | *Lactobacillus* | *L. johnsonii* |
| 0.026 | Bacteria | *Firmicutes* | *Bacilli* | *Lactobacillales* | *Streptococcaceae* | *Streptococcus* | *S. respiraculi* |
| 0.029 | Bacteria | *Firmicutes* | *Bacilli* | *Lactobacillales* | *Lactobacillaceae* | *Lactobacillus* | *L. murinus* |
| 0.037 | Bacteria | *Firmicutes* | *Bacilli* | *Lactobacillales* | *Streptococcaceae* | *Streptococcus* | *S. suis* |
| 0.043 | Bacteria | *Firmicutes* | *Bacilli* | *Lactobacillales* | *Lactobacillaceae* | *Lactobacillus* | *L. reuteri* |

Seven of the 11 species derived from the *Lactobacillales* order, three belonging to *Enterobacterales* and the remaining one species in the order *Chromatiales*. The *Lactobacillales* order was further divided into four beneficial *Lactobacillus* and three animal-related *Streptococcus* spp. (Table 2). The relative abundance of *L. Murinus* was much higher than any other 10 age-related species in the newborn, adolescent, and adult stages, but significantly decreased in the seniors compared to the newborn and adult rats (Figure 2c; pairwise Wilcoxon test, adjusted p < 0.05, Table S3). Previous animal studies have shown the protective effects of commensal bacteria *L. murinus* against gastrointestinal diseases [41,42] and viral respiratory infections [43]. We connected a possible role of *L. murinus* in rat normal development and aging processes through bioinformatics analyses, although further experimental validations are needed. Among the selected 11 species, the relative abundance of *Salmonella enterica*, which is a high prevalence species of the *Enterobacterales* order, was significantly increased in the elderly compared to the adolescent rats (Figure 2c; pairwise Wilcoxon test, adjusted p < 0.05, Table S3). Although *S. enterica* is an enteric pathogen of both humans and animals, it can spread to all tissues and rodents are natural reservoirs of *Salmonella* [44]. Perhaps *Salmonella* infection and spreading is more common in older rats where their immune functions tend to decline with age. Other age-dependent species that were highly enriched in elderly rats include *S. suis*, *S. himalayensis*, and two environmental species (*A. prosperus* and *Pantoea* sp. At-9b) (Table S3). In addition, the abundances of *L. reuteri, L.johnsonii,* and *Candidatus Arsenophonus lipoptenae* were much higher in adolescent rats than in other stages (Table S3). Adolescent rats also showed the significant lowest microbial alpha diversity at the order and species levels compared with the newborn and adult rats (Figure 2e-f; pairwise Wilcoxon test, adjusted p < 0.05), although with no significant beta diversity differences compared to the other three developmental stages (Figure 2g,h,i).

### 3.3 Lung microbial composition changes dramatically during the four developmental stages

The above 11 stage-related species were identified at the whole/bulk tissue level, which may not capture the complexity of the tissue-specific microbial changes during development. Therefore, we first processed the operational taxonomic unit (OTU) table at each developmental stage individually to compare the inter-tissue microbial composition changes in this section, and in section 3.5 to identify and compare the age-dependent species for each tissue type. *Bacillales* was prominent with the relative abundance over 40% in order among all tissue groups (Figure 3a,d,g,j; Table S4). Liver and muscle generally have the lowest microbial alpha diversity as compared to the other organs across the 4 developmental stages (Figure 3b,e,h,k; pairwise Wilcoxon test, adjusted p < 0.05).

**Figure 3.**
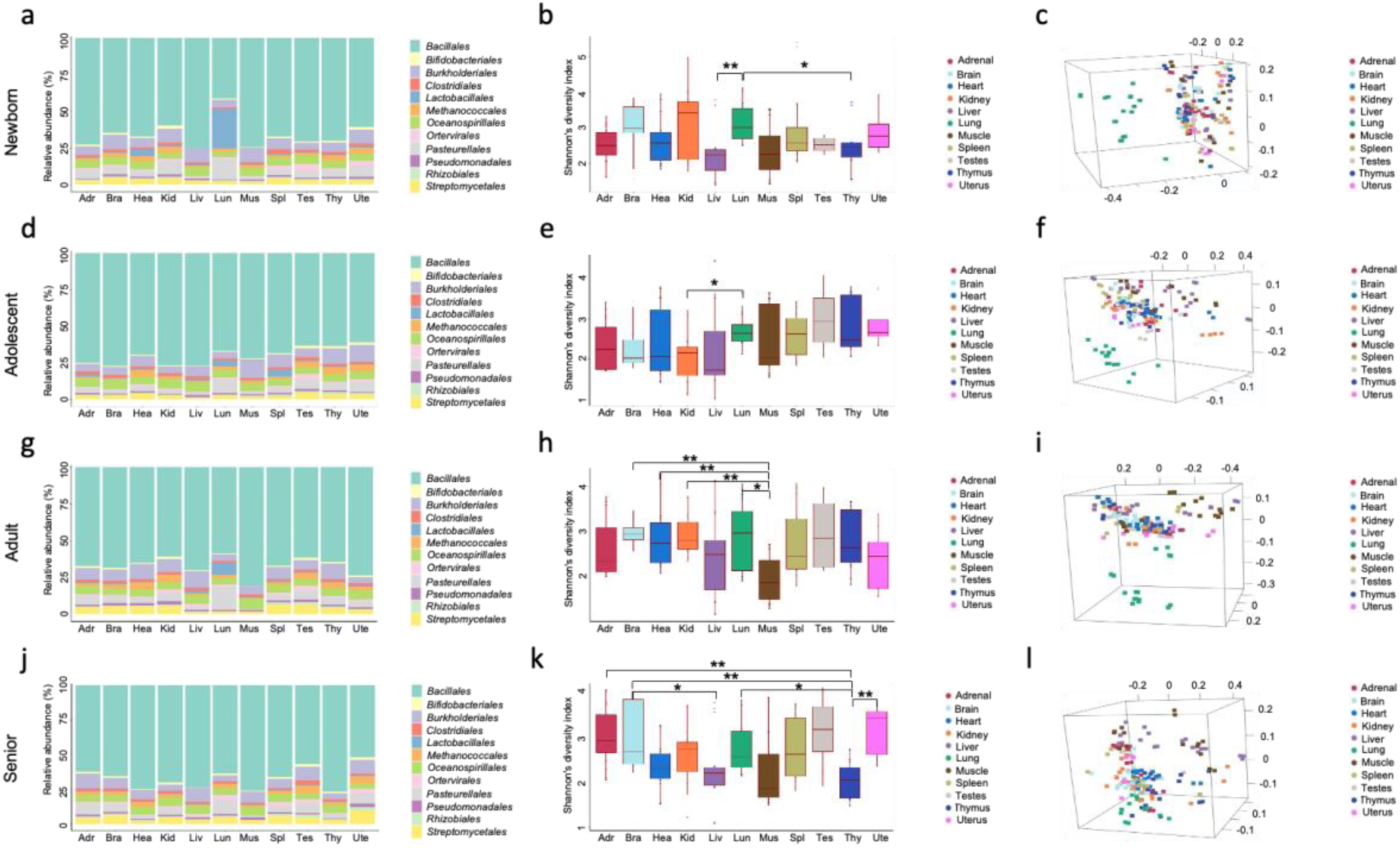
Tissue-level microbial composition and diversity comparisons during the four developmental stages. Taxa composition bar plots illustrate the microbial relative abundance (%; in y-axis) of the top 12 most abundant orders at the newborn (a), adolescent (d), adult (g), and senior stage (j). Species-level Shannon’s diversity index (in y-axis) of the 11 tissues at the newborn (b), adolescent (e), adult (h), and senior stage (k). Pairwise Wilcoxon rank sum tests with BH correction were used to test for diversity differences between tissues. Only statistically significant comparisons (p < 0.05) were marked with a single star (*), and p-values < 0.01 were marked with two stars (**). 3D-view species-level PCoA of Bray-Curtis dissimilarity between samples colored in 11 different colors according to each tissue type at the newborn (c), adolescent (f), adult (i), and senior stage (l).

In newborn rat samples (n=168), pairwise comparisons showed significantly higher *Lactobacillales* abundance in lungs compared to other 10 tissues (Figure 3a; adjusted p < 0.05, Table S4-5). The heart has the second highest relative abundance of *Lactobacillales* although not statistically significant compared with the remaining organs (Figure 3a; Table S4-5). Lung samples also have the highest abundance of *Pasteurellales* in newborn rats, and the pairwise comparisons were all significant except for that with kidney (Figure 3a; adjusted p < 0.05, Table S4-5).

A significant decrease of *Lactobacillales* in lungs was observed between newborn and adolescent rat samples (Figure 3a,d; adjusted p < 0.05, Figure S1a). The spleen has a slightly higher relative abundance of *Lactobacillales* than that of the lung at the adolescent stage (Figure 3d; Table S4). Other order-level enrichments in adolescent rats include: the muscle and uterus samples have the first and second highest abundances of *Burkholderiales*, respectively (Figure 3d; Table S4). Except for the testes and uterus samples, the lung has a significantly higher abundance of *Pasteurellales* than other tissues (Figure 3d; pairwise Wilcoxon test, adjusted p < 0.05, Table S4-5).

*Pasteurellales* remains to be the most predominant bacteria order in lung tissues compared to any other tissues of adult (n=163) and senior rats (n=167) (Figure 3d,g,j; Table S4). A slight increase of the relative abundance of *Lactobacillales* was detected in adult lungs compared to adolescent lungs (adjusted p = 0.207; Figure S1a). Similar with newborn lungs, adult lungs have significantly higher *Lactobacillales* abundance compared to other 10 tissues (Figure 3d,g; pairwise Wilcoxon test, adjusted p < 0.05, Table S5). In addition, the relative abundances of *Streptomycetales* in the liver, muscle, lung, and uterus were much lower than in other tissues in adult rats (Figure 3g; Table S4).

The relative abundances of *Lactobacillales* in elderly samples were no longer significant between the lung and other tissues; actually their abundances were under detectable levels across all tissue types (0.21%-0.67%) (Figure 3j; Table S4-5). Moreover, the lung microbiomes showed distinct separation from other tissues and shifted significantly in elderly rats compared with younger animals (Figure 3c,f,i,l). These observations not only suggested that the lung microbiome was more unique and vulnerable at older age, but also indicated the existence of a potential positive correlation between the abundance of *Lactobacillales* with the development of the lung microbiota.

### 3.4 *Lactobacillus* and *Bifidobacterium* were detected by PCR in healthy lungs

Lung microbial compositions, particularly those of *Lactobacillales,* were more likely to decrease as rats grow older, emphasizing the importance of maintaining the amount of “good” bacteria in lungs. In order to verify that *Lactobacillus* are present in healthy lungs, both conventional and real-time PCR were performed in adult rat and human lung tissues. *Bifidobacterium* was also selected as the two bacteria genera exhibited similar beneficial effects [45]. Genus-specific PCR primers which target a total of 15 and 30 *Lactobacillus* and *Bifidobacterium* spp., respectively, were taken from [25] and used for the detection and quantification of the corresponding bacteria in our study.

Different from that observed in *Lactobacillales* (Figure S1a), no significant abundance changes were found for *Bifidobacteriales* in lungs among the four stages in the discovery dataset (Figure S1b). The average abundance of *Lactobacillales* was slightly higher than that of *Bifidobacteriales* in both discovery and validation dataset 1 (Figure S1c-d; not reaching the statistical significance), which were based on RNA-Seq-derived adult rats’ microbial results. Similarly, RT-PCR detected positive for *Lactobacillus* and *Bifidobacterium* in all 6 rat lung samples with no statistical abundance differences from the validation dataset 1 (Table 3; Figure S1e). Furthermore, up to 15 and 30 *Lactobacillus and Bifidobacterium* spp., respectively, were detected in healthy human lungs, and *Lactobacillus* bands were generally brighter and more unique than that of *Bifidobacterium* (Figure S1f). Subject A11 with the highest BMI, tested negative for *Lactobacillus,* and has been enriched with different *Bifidobacterium* spp. compared with the other two subjects (Figure S1f).

**Table 3.** Summary of the quantifying microbial abundances by RT-PCR.

| | Sample | *Repli1<br>Ct | *Repli2<br>Ct | *Repli3<br>Ct | Mean<br>Ct | ΔCt | $\Delta\Delta$Ct | $2^{-\Delta\Delta\text{Ct}}$ |
| --- | --- | --- | --- | --- | --- | --- | --- | --- |
| ***Actb***<br>**(house-keeping)** | C1123 | 13.569 | 13.339 | 13.873 | 13.594 | -- | -- | -- |
|  | C1450 | 12.959 | 12.977 | 12.951 | 12.962 | -- | -- | -- |
|  | C1456 | 14.024 | 13.694 | 13.873 | 13.864 | -- | -- | -- |
|  | C1151 | 11.741 | 14.112 | 13.704 | 13.186 | -- | -- | -- |
|  | C1451 | 13.009 | 13.022 | 13.046 | 13.026 | -- | -- | -- |
|  | C1460 | 13.802 | 13.803 | 13.887 | 13.831 | -- | -- | -- |
| ***Lactobacillus* spp.** | C1123 | 33.859 | 32.921 | 31.866 | 32.882 | 19.288 | 0.440 | 0.737 |
|  | C1450 | 32.307 | 30.922 | 31.172 | 31.467 | 18.505 | -0.343 | 1.268 |
|  | C1456 | 29.119 | 29.097 | 29.805 | 29.340 | 15.476 | -3.372 | 10.351 |
|  | C1151 | 32.109 | 31.662 | 34.142 | 32.637 | 19.451 | 0.603 | 0.658 |
|  | C1451 | 30.818 | 31.421 | 32.480 | 31.573 | 18.547 | -0.301 | 1.232 |
|  | C1460 | 37.193 | 34.168 | 33.787 | 35.049 | 21.218 | 2.370 | 0.193 |
| ***Bifidobacterium* spp.**<br>**(reference)** | C1123 | 32.284 | 32.872 | 32.728 | 32.628 | 19.034 | 0.186 | 0.879 |
|  | C1450 | 32.381 | 31.848 | 32.597 | 32.275 | 19.313 | 0.465 | 0.724 |
|  | C1456 | 31.133 | 31.566 | 32.459 | 31.719 | 17.855 | -0.993 | 1.990 |
|  | C1151 | 32.652 | 32.867 | 32.980 | 32.833 | 19.647 | 0.799 | 0.574 |
|  | C1451 | 31.429 | 32.678 | 31.910 | 32.006 | 18.980 | 0.132 | 0.912 |
|  | C1460 | 31.568 | 32.330 | 32.366 | 32.088 | 18.257 | -0.591 | 1.506 |
|  | Reference average | -- | -- | -- | -- | 18.847 | -- | -- |

\*Repli: Replicate

### 3.5 Comparison of age-dependent microbial species among different tissues

Kruskal’s tests adjusted for multiple comparisons at the species level were then carried out to identify the age-dependent microbes for each tissue type, respectively. Only the lung, testes, thymus, kidney, adrenal, and muscle were found to have one to 52 age-dependent species (adjusted p < 0.05, Table S6).

There were a total of 52 species, of which 32 derived from the order of *Lactobacillales,* together with seven *Bacillales* spp., four *Ortervirales* spp., two *Campylobacterales* spp., and seven other species from seven different bacteria orders had been identified to be the age-dependent species in lungs (Figure 4a; adjusted p < 0.05, Table S6). The *Lactobacillales*, *Ortervirales, Campylobacterales,* and *Bacillales* were found to be the top 4 abundant orders associated with lung development and aging (Figure 4a). The loss of *Lactobacillales* in elderly lungs (Figure 4a) agrees well with our previous taxonomy investigation at the order level (Figure 3a,d,g,j). Only the lung samples have 8 overlapped species with the stage-related microbes identified at the whole/bulk tissue level, and 6 of them were from the order of *Lactobacillales* (Figure 4g; Table S6), further indicating there are strong associations between *Lactobacillales* spp. with lung microbiota development and maturation.

**Figure 4.**
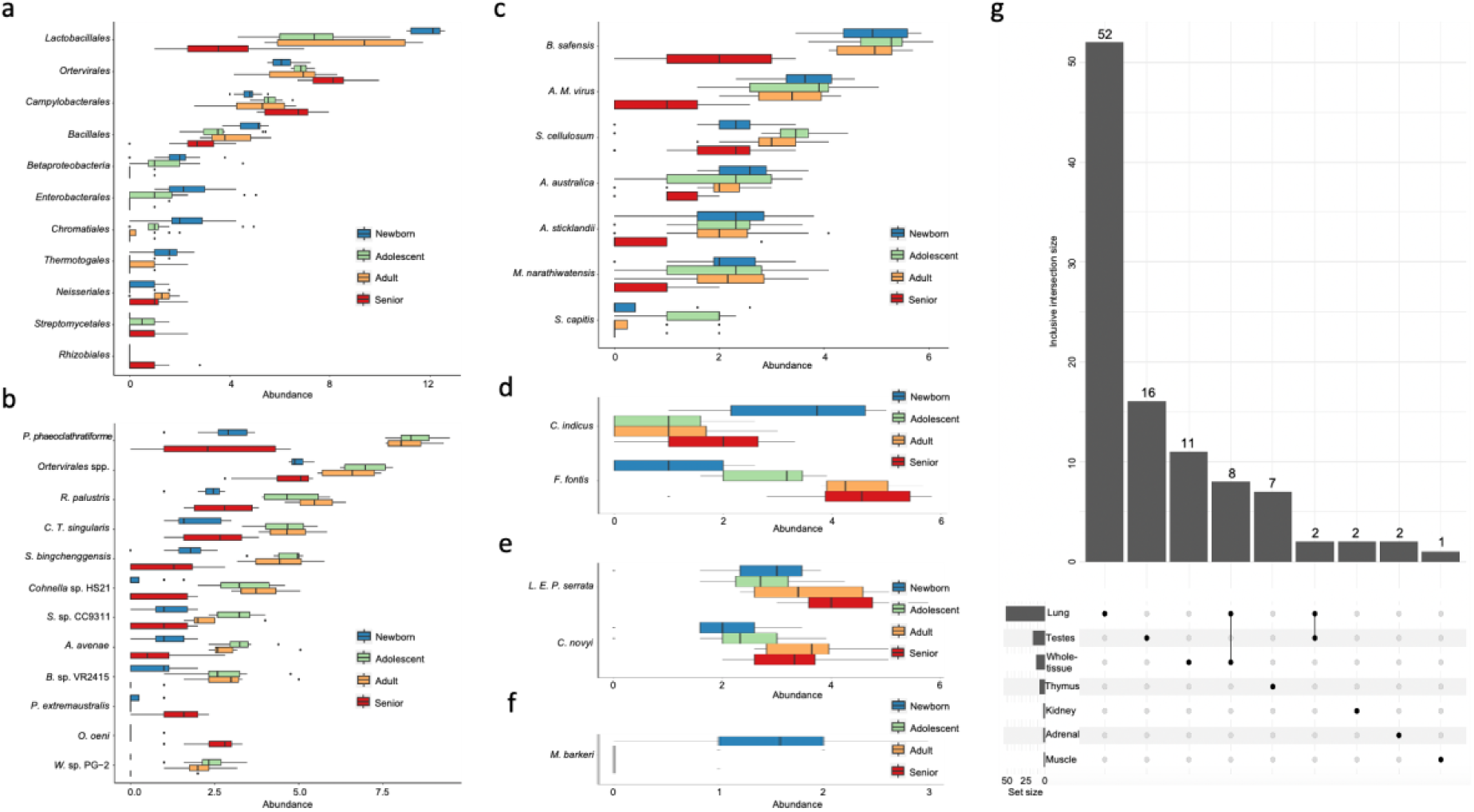
Comparison of age-dependent microbial species among six different tissues. Microbial abundances of the selected microbes (in x-axis) of the four stages in lung (a), testes (b), thymus (c), kidney (d), adrenal (e), and muscle (f) samples. An UpSet plot showing the intersected stage-related microbial species in the 6 tissues and whole-tissue levels (g). Each bar in the bar chart shows a different combination of tissues and the size of inclusive intersected species (values in y-axis). The graphical table below the bar chart indicates the corresponding memberships. Each row in the table is one of the 6 tissues or whole tissue. The filled black dots and lines show the combination of tissues that have sets intersections. A smaller bar chart on the left side of the graphical table displays the size of elements per set.

There were 5 animal-derived *Ortervirales* and 11 environmental species identified to be the age-dependent microbes in testes (Figure 4b; adjusted p < 0.05, Table S6). Among them, only *P. extremaustralis* and *O. oeni* were significantly enriched in the senior testes samples (Figure 4b); and the remaining species were significantly enriched in the adolescent and adult samples (Figure 4b). Two murine leukemia viruses, namely *Murine leukemia virus* and *Murine leukemia-related retroviruses*, were commonly identified from the lung and testis tissues (Table S6).

In thymus samples, we found a total of seven species that changed their abundances during the whole development process (Figure 4c; adjusted p < 0.05, Table S6). Among them, there were five environmental and one animal virus (*B. safensis, A. australica, A. sticklandii, M. narathiwatensis, S. cellulosum*, and *Avian myeloblastosis virus*) decreased in elderly samples (Figure 4c). In addition, *S. capitis*, a commensal skin microbe was significantly enriched in the adolescent samples (Figure 4c).

Lastly, there were five species that may be involved in the developmental processes in the kidney, adrenal, and muscle samples, respectively. More specifically, the significant increase and decrease of two environmental microbes in the kidney, namely *C. indicus* and *F. fontis* were found in newborn samples, respectively (Figure 4d). Two pathogens namely *Legionella endosymbiont of Polyplax serrata* and *Clostridium novyi* were significantly enriched between the older (adult + elderly) vs young (juvenile + adolescent) in the adrenal samples (Figure 4e). *Methanosarcina barkeri*, a methanogenic archaea was found to be significantly decreased in the adolescent, adult, and senior stages compared to newborn muscle samples (Figure 4f).

### 3.6 Identification of four rat inter-tissue microbial heterogeneity patterns

As no notable batch effects in PcoA were observed over the four course of aging (Figure 3c,f,i,l), we next performed the unsupervised cNMF [30] to assess the inter-tissue microbial heterogeneity regardless of the ages of the rats involved. The 2829 species-level OTU abundance table of all rat samples (n=660) was used as input for cNMF analysis. cNMF identified a total of four inter-tissue microbial heterogeneity patterns (P1-P4), in which the cluster number K=4 corresponds to the maximum stability in the data (Figure 5a). The local density filtering threshold was set at ∼0.2 based on KNN imputation and the consensus clustergram (Figure 5b-c). 357 microbial species were subsequently identified as the signatures (or called meta-microbe) for each of the four patterns, respectively (Tables S7). The microbial abundance heatmap with samples and species ordered according to their predicted pattern, showed four well separated microbial heterogeneity patterns (Figure 5d).

**Figure 5.**
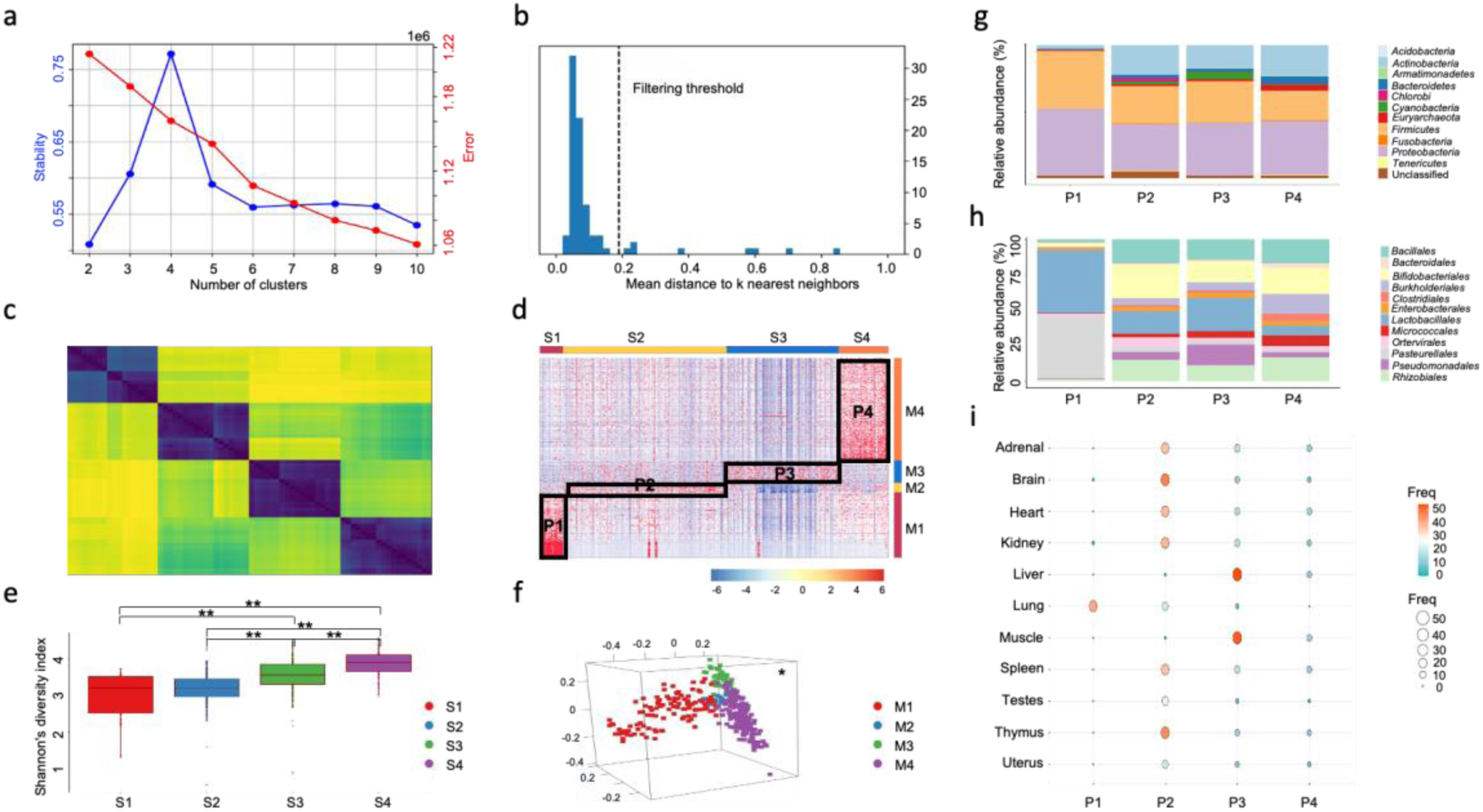
Identification of four rat inter-tissue microbial heterogeneity patterns. (a). The K selection plot by using the trade-off between solution stability (primary y-axis; in blue color) and solution error (secondary y-axis; in red color). K = 4 was selected as the optimal number of clusters as it is highest in stability and has shown relatively lower error rate. (b). Set an outlier threshold of ∼0.2 based on inspecting the histogram of distances between each cluster and its K nearest neighbors. (c). Clustergram diagnostic heatmap of the chosen K = 4 showed that there is a high degree of agreement between the replicates with a few outliers. (d). The microbial abundance heatmap with 660 samples in columns (S1-S4) and 357 species in rows (M1-M4), ordered according to their predicted cluster memberships, showed four well separated microbial heterogeneity patterns (P1-P4). In the heatmap, red indicates higher abundances and blue represents lower abundances. (e). 357 microbial species-level Shannon’s diversity index (in y-axis) of the four patterns. Pairwise Wilcoxon rank sum tests with BH correction were used to test for diversity differences between patterns. Only statistically significant comparisons (p < 0.05) were marked with a single star (*), and p-values < 0.01 were marked with two stars (**). (f). 3D-view 357 species-level PCoA of Bray-Curtis dissimilarity between microbial communities colored in 4 different colors according to each community of microbes. Pairwise multilevel comparisons using adonis with the Bonferroni correction were used to test for beta diversity differences between communities. All pairwise comparisons were significant (p < 0.05), and were marked with a single star (*) at the top right position. Taxa composition bar plots illustrate the microbial relative abundance (%; in y-axis) of the top 12 most abundant microbes at the phylum (g) and the order level (h) based on the 357 pattern-specific species’ microbial profile. (i). A balloon plot to summarize and compare the tissue distributions (rows) for each pattern (columns), where each cell contains a dot whose color and size reflect the magnitude of a numerical value (Freq).

The 660 rat samples were classified into four inter-tissue subtypes (S1-S4; Tables S7). S4 had the highest microbial alpha diversity, followed by S3, and the lowest in S1 and S2. Although no significant alpha diversity differences were found between S1 and S2, all other pairwise alpha diversity comparisons between subtypes were significant (Figure 5e; pairwise Wilcoxon test, adjusted p < 0.05). The 357 subtype-specific microbial signatures include 71 different orders belonging to 18 different phyla (Tables S7). *Proteobacteria*, *Firmicutes*, *Actinobacteria*, *Viral*, and *Bacteroidetes* were the top five major phyla (Figure 5g). 117, 17, 40, and 183 microbial signatures for each tissue subtype were used to define the four microbial communities (M1-M4). All pairwise beta diversity comparisons among the communities were significant (Figure 5f; pairwise PERMANOVA, adjusted p < 0.05).

The four inter-tissue microbial patterns (P1-P4) each consist of the corresponding tissue subtypes (S1-S4) and microbial communities (M1-M4). S2 contains the largest number of rat samples (310), spanning all 11 different tissue types, but with only one liver and one muscle sample (Figure 5i; Table 4). 17 S2 specific microbial species were present in M2, with significant higher relative abundances of *Bifidobacteriales* and *Ortervirales* compared with the other microbial communities (Figure 5h). Liver and muscle samples were almost exclusively grouped into S3 and S4 (Figure 5i; Table 4). Similar to S2, S3 has a large sample size (212) with all 11 tissues. A total of 40 bacteria species were grouped into the M3, and were highly enriched in the 212 samples from S3 (Figure 5d). The relative abundances of *Pseudomonadales* and *Enterobacterales* in P3 (consist of 212 samples and 40 species) were much higher than other patterns (Figure 5d). S4 includes 94 samples but with no lung tissue (Figure 5i). P4 has higher *Burkholderiales*, *Micrococcales*, and *Clostridiales*, and lower *Lactobacillales* relative abundances (Figure 5h). S1 is the smallest subtype with 44 samples, and 41 of them come from the lung (Table 4; Figure 5i). *Lactobacillales* and *Pasteurellales* were the two most dominant microbial signatures in M1 (Figure 5h).

**Table 4.**
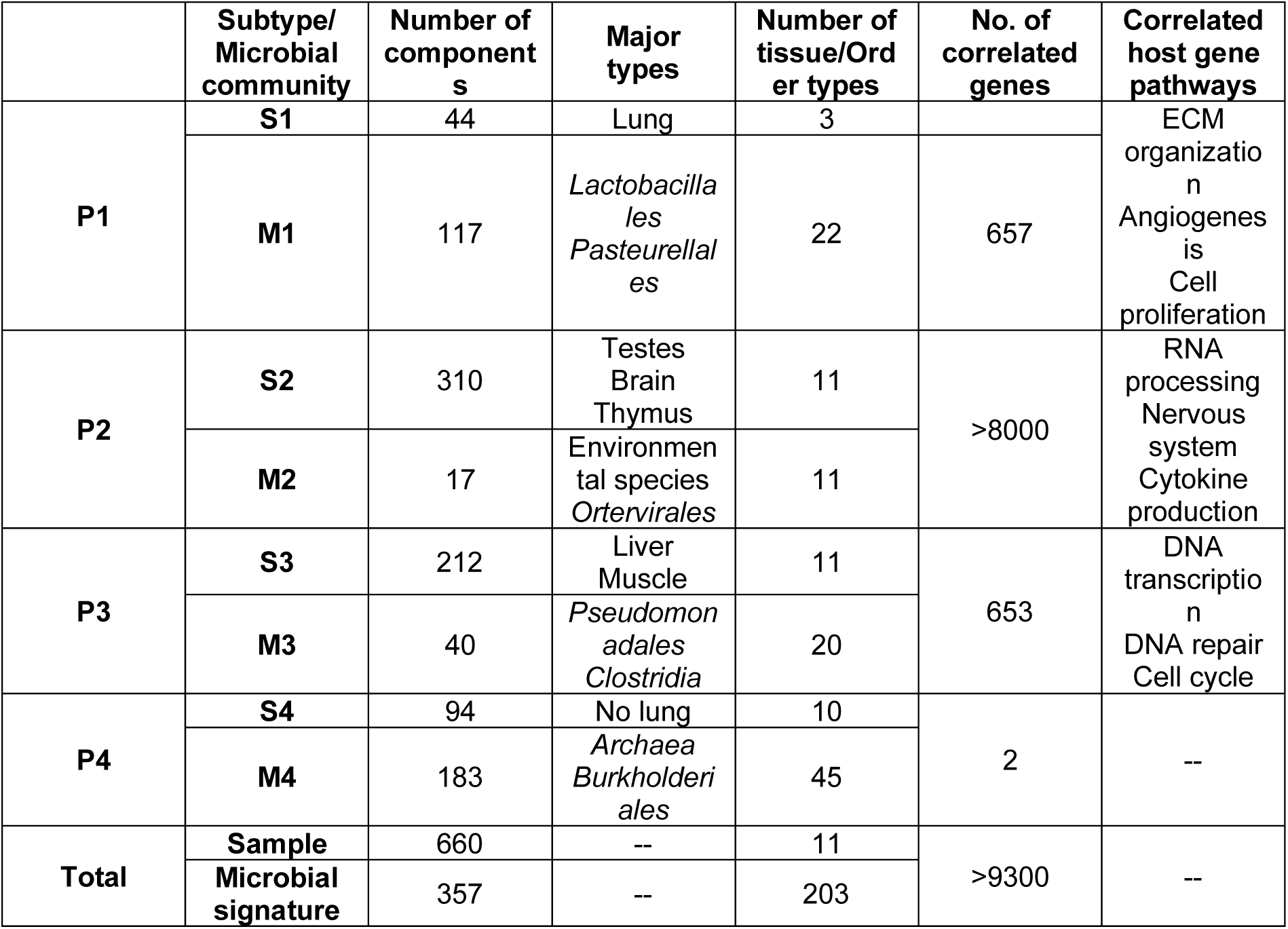
The distributions and characters of the inter-tissue subtypes and microbial signatures.

### 3.7 Pattern-specific microbe-host gene interactions

We inferred a microbe-host gene interaction network individually for the 4 inter-tissue microbial patterns identified in the study. Spearman’s correlation analyses were used to predict putative interactions between microbial species abundances and host gene expressions in each pattern.

After filtering, a total of 673 genes were found significantly positive/negative correlated with the abundances of the 117 microbial signatures in P1 (adjusted p < 0.05, Table S8). The genes and microbes were organized by hierarchical clustering, and their Spearman’s correlation coefficients were visualized in a heatmap (Figure 6a). The absolute correlation coefficients less than 0.3 were marked as zero to assist better visualization. Although most of the genes appear to be lowly correlated with the majority of microbial signatures, four notable enrichments were depicted in the microbe-host gene heatmap (Figure 6a). Of note, the correlations with genes reflect their combination, rather than the species themselves. More specifically, the largest cluster C2 contained more than 79% (n=534) of the selected 673 genes which were either positively or negatively correlated with 41 microbial species (Figure 6a, Table S8). The 15 *Streptococcus* spp. and five *Lactobacillus* spp. derived from the order of *Lactobacillales*, and four other species (namely *Candidatus Arsenophonus lipoptenae, Acidihalobacter prosperus, bacterium* 2013Arg42i, and *Bacillus coagulans*) as a whole community was positively correlated with these 534 genes (Figure 6a; adjusted p < 0.05, Table S8). Meanwhile, two viruses (*Murine leukemia virus* and *Spleen focus-forming virus*), *S. enterica*, and the other 14 bacteria were negatively correlated with those host genes. This largest set of genes is significantly involved in extracellular matrix (ECM) organization, angiogenesis, cell proliferation, and Wnt signaling pathways (adjusted p < 0.05, Table S8), which might be co-regulated by the microbes and hosts. The remaining three clusters of genes were associated with bacteria infection (C4), glucose transporters (C3), and T cell activation (C1, not significant) pathways and processes (Table S8).

**Figure 6.**
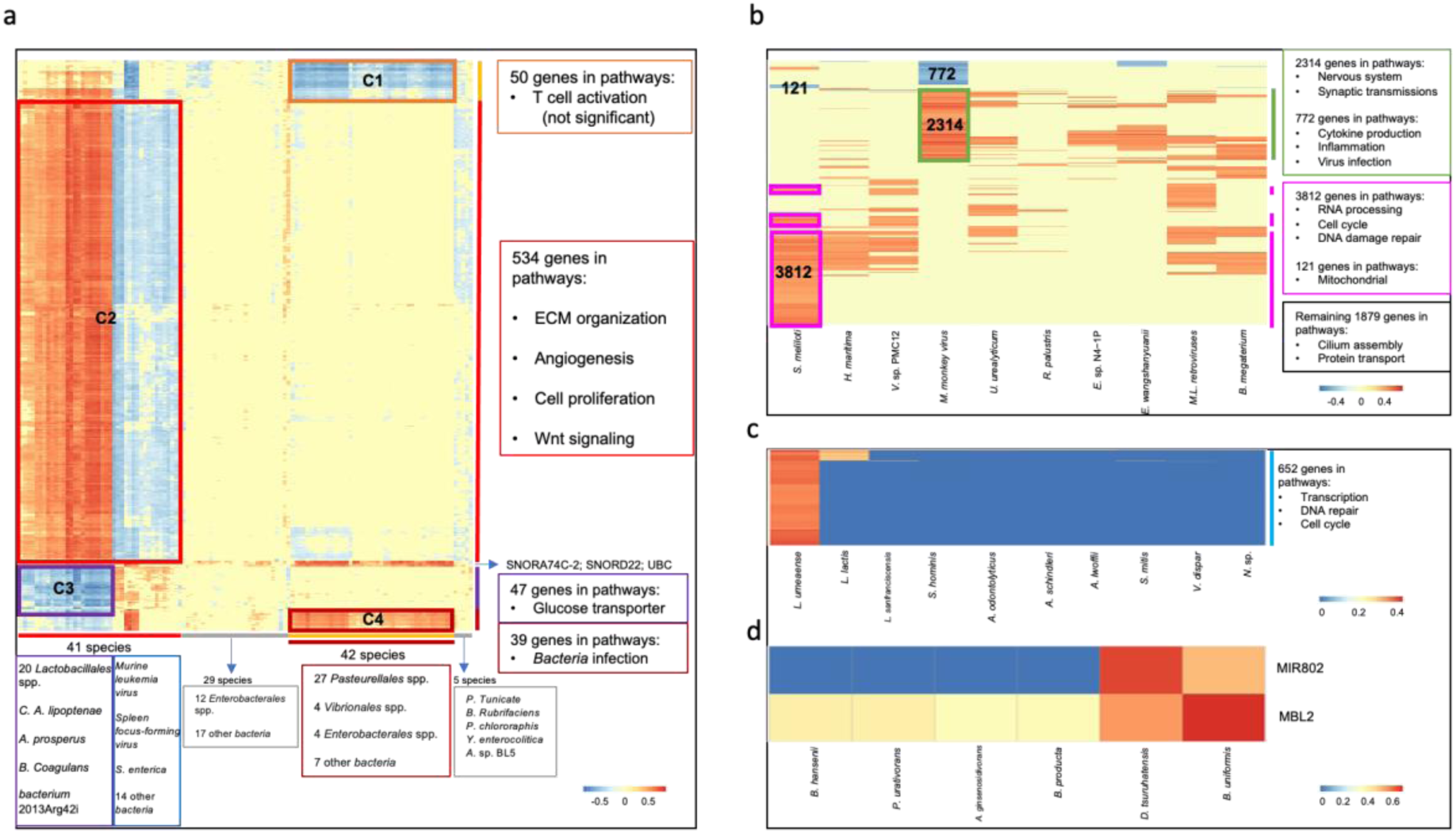
Pattern-specific microbe-host gene interactions. Heatmaps of the Spearman’s correlation coefficients between the pattern-specific species (in columns) with the host genes (in rows and in human gene symbols) in P1 (a), P2 (b), P3 (c), and P4 (d). Red indicates positive correlations and blue represents negative or low-positive correlations. The genes and species were hierarchically clustered (complete linkage, Euclidean distance), and their associated characteristics (such as taxonomic ranks and pathway categories; if any) were shown on the right hand of the corresponding heatmap, respectively. Gene cluster memberships (C1-C4) were shown in (a) and gene cluster numbers were shown in (b).

The 17 microbial species in P2 were significantly associated with a sum of 9371 host genes (adjusted p < 0.05), while around 40% of these microbes each correlated with less than 200 genes. We filtered out microbes with fewer than 200 correlated genes, and the 0.3 absolute correlation coefficient cutoff was applied to remove lowly correlated genes, resulting in a microbe-host gene heatmap with 10 species and 8780 host genes (Figure 6b; Table S8). The heatmap showed that 3812 and 2314 genes were significantly positively, while 121 and 772 genes were significantly negatively correlated with the abundances of *S. meliloti* and *Mason-Pfizer monkey virus*, respectively (Figure 6b; adjusted p < 0.05, Table S8). RNA processing, cell cycle, DNA damage repair related pathways were significantly overrepresented in the 3812 genes which were positively correlated with *S. meliloti*. The 121 *S. meliloti* negatively correlated genes were enriched in mitochondrial related pathways (adjusted p < 0.05, Table S8). *Mason-Pfizer monkey virus* appears to be positively associated with 2314 host genes in pathways of nervous system and synaptic transmissions. The 772 negatively correlated genes involved in various cytokine production, inflammation, and virus infection related pathways (adjusted p < 0.05, Table S8). The remaining positively correlated 1879 genes, which were partially shared between microbes including *Murine leukemia-related retroviruses*, were enriched with cilium assembly, cilium movement, and protein transport related pathways (Table S8). Cilium related genes and pathways were also positively correlated with the abundance of *Exiguobacterium* sp. N4-1P.

Gut-derived bacteria *Lachnoanaerobaculum umeaense* was the most significantly positive associated with 652 host genes in P3 (Figure 6c; adjusted p < 0.05, Table S8). KEGG and GO enrichment analysis indicated that these genes were involved in pathways such as transcription, DNA repair, and cell cycle (adjusted p < 0.05, Table S8). RNA binding gene *RBM12B* was positively correlated with another five bacteria species in P3 including *Lactobacillus sanfranciscensis*, *Acinetobacter lwoffii*, *Acinetobacter schindleri*, *Actinomyces odontolyticus*, and *Staphylococcus hominis* (adjusted p < 0.05, Table S8). There were 72 genes that had correlations above 0.3 with the abundance of *Lactococcus lactis,* and enriched for cell cycle related pathways (adjusted p < 0.05, Table S8). Lastly, only the soil bacteria *Nocardioides* sp. MMS17-SY207-3 was found to be positively correlated with the *TMCC1* gene with unknown function.

Among the 183 microbial signatures in P4, only *Bacteroides uniformis* and pathogen *Delftia tsuruhatensis* each had significant positive correlations with host genes (Figure 6d; adjusted p < 0.05, Table S8). More specifically, *MBL2*, which is a component of innate immunity [46], has been found to be positively correlated with the abundance of *B. uniformis* and *D. tsuruhatensis* in our study. *MIR802,* a glucose metabolism regulator [47], is more positively correlated to an increase in *D. tsuruhatensis*’s abundance.

## 4. Discussion

Rats are commonly used in laboratory settings where little information is available about the composition and diversity of their microbial communities. In our study, we systematically investigated the spatial and longitudinal microbial patterns in various healthy rat body compartments and in different developmental stages. Lung microbial composition changes dramatically, and liver with muscle exhibits the lowest microbial alpha diversity as compared to other organ systems across the 4 major life-history stages of rats. The lung, testes, thymus, kidney, adrenal, and muscle microbial habitats were found to have one to 52 species that changed their abundances as the rats got older. Moreover, potential positive correlations between the abundance of *Lactobacillales* and lung maturation to lung aging are inferred from our study.

Species in the order of *Lactobacillales* are commonly called lactic acid bacteria (LAB). LAB are a group of gram-positive and lactose fermenters, which have long been known for their health-promoting properties of milk. Previous studies showed that microbial colonization of mucosal intestinal tissues during the period of suckling shapes the infant’s immune system [14]. Accumulating evidence indicates that at this stage microbes (i.e. LAB and *Bifidobacterium*) are transmitted from mother to offspring mainly through breast milk [14,48,49]. In addition to fundamental nutrients and bioactive compounds, human breast milk features a unique microbiome, including mutualist, commensal, and probiotic potential bacteria species [50]. The Food and Agriculture Organization of the United Nations and the WHO (FAO/WHO) defines probiotics as “live microorganisms which when administered in adequate amounts confer a health benefit on the host.” [51]. Thus, previous studies established a link between immune-enhancing properties and LAB in the gut. In our current study, we found members of LAB (i.e. *L. murinus* and *L. animalis*) which show significant overlap with known immune-related LAB, were positively correlated with a large number of host genes involved in cell migration and proliferation, especially in the lungs. A gradual decline of lung LAB abundance was strongly associated with older age, which relates to a metabolic process, suggesting that LAB could be potentially used as an anti-aging probiotics.

Following this finding, we aim to identify more microbial signatures that reflect the inter-tissue microbial heterogeneity irrespective of the developmental stages, in this spatially and temporally heterogeneous data. When compared with core species, uncommon and rare microbes are less observed, but their overall numbers are abundant with important ecological roles individually or in groups [52]. We previously used a microbial prevalence threshold at 1% to eliminate rare and potentially contaminated species in two human-associated microbiome studies [26,53]. Given the fact that rodent microbiomes are more likely to be contaminated with fecal matter, and more than two technical replicates are available in the discovery dataset, we increased the threshold to 10% to mitigate the effects of contamination. Based on the identified 2829 species as global-level microbiome constituents, four inter-tissue microbial heterogeneity patterns (P1-P4) were identified in an unsupervised fashion, with a total of 357 pattern-specific microbial signatures to distinguish among different patterns. The 357 microbial signatures come from 71 different orders in 18 different phyla. *Proteobacteria*, *Firmicutes*, *Actinobacteria*, *Viral*, and *Bacteroidetes* were the top five most abundant phyla. There were 12 *Halobacteria* in the 357 microbial signatures, which are members of the domain Archaea and are capable of surviving in both low and high salt environments [54]. Using culture-dependent and - independent techniques, various halophilic archaea have been identified in salted food products like fish sauce and table olives [55]. Moreover, the existence of enteric halophilic archaea in the human microbiome is now generally accepted [56,57]. Thus, it is likely that the *Halobacteria* spp. we identified are food-borne microbial species in rats. The 12 *Halobacteria* together with another 2 Archaea species, namely *Methanocaldococcus infernus* and *Candidatus Nitrosocosmicus franklandus,* were microbial signatures that all come from the P4. Four *Ortervirales* spp. plus one *Leucania separata nucleopolyhedrovirus* from P1 and P2, are the only 5 viral species in the 357 signatures. Microbial signatures in each pattern were selected to investigate the potential microbe-host gene interactions. Since *Archaea* species account for smaller proportions, and have fewer correlations with host genes compared to the remaining signatures. Therefore, we focused our attention on the composition and diversity of 338 bacteria and five viral species in each pattern.

More than 93% of the cases in P1 were lung samples. The relative proportion of order *Lactobacillales* in *Firmicutes* and *Pasteurellales* in *Proteobacteria* in P1 account for more than 85% of all 357 microbial species. The relative abundance of *Actinobacteria* was significantly decreased in P1 compared with the remaining three patterns. P2 and P3 contained all 11 tissue types, and no lung samples were found in P4. P2 was the largest pattern representing 41.1% (n=310) of all rat samples, but only 17 species were considered as the markers for this pattern. The relative abundances of *Bifidobacteriales* and *Ortervirales* in P2 were much higher than the other patterns. *Pseudomonadales* and *Enterobacterales* were significantly increased in P3. P4 has higher *Burkholderiales*, *Micrococcales*, and *Clostridiales*. Microbial signatures in each pattern were mostly positively correlated with different host genes in different metabolic and biological functions. For example, 20 LAB and four other species in P1 were correlated with genes involved in ECM organization, cell migration and proliferation signaling pathways. *L. umeaense* in P3 was positively associated with DNA transcription and cell cycle. *B. uniformis* in P4 with limited innate immune signaling association. The signatures in P2 largely come from environmental sources such as soil and water sediment. Most environmental species are free-living and widely distributed in multiple habitats, which are capable of colonizing and/or causing infections in mammals [58–60]. Compared to germ-free rats, conventional rats are exposed to enriched environments. Thus, it is not surprising that there are 10 environmental microorganisms within the 19 core species identified in the study, of particular interest are *S. meliloti* and *Exiguobacterium* sp. N4-1P. They are two of the microbial signatures in P2, which showed higher abundances in the P2 host samples and lower abundances in most liver and muscle tissues. The majority of the correlated host genes and pathways for the two species in P2 samples were positive. For example, *S. meliloti* has been significantly positively correlated with thousands of genes involved in RNA processing and DNA damage repair. Cilium related genes and signaling pathways were positively correlated with the abundance of *Exiguobacterium* sp. N4-1P. Therefore, species from environmental sources contribute to host microbial diversity and strongly interact with host genes in various cellular functions.

Our study has limitations and the largest was the lack of investigating the paired GI samples. The GI microbiome is among the most studied and best characterized microbial niche to date. GI microbiota are distinct between wild and laboratory-kept rodents [61]. In laboratory settings, a wide variety of symbiotic microorganisms harbor in the rat digestive tract and it is primarily composed of *Firmicutes* and *Bacteroidetes* phyla [62]. 16S amplicon sequencing and RT-PCR-based studies have shown that LAB (within the *Firmicutes* phylum) were the most abundant bacteria in the rat gut microbiota [63–65]. What’s more, LAB induces a protective effect, and is one of the most predominant microbes in the feces of the breast-fed neonatal rats [19]. Laboratory rodent diets can be a major source of microbial communities in the GI tract [66]. Rats are omnivorous, eating a variety of plant and animal food items. Grain-based and purified diets represent two of the major commercially made diets being used in laboratory rodent studies [66,67]. Grain-based diets contain cereal grains, wheat middlings, animal byproducts, many non-nutrients, and contaminants, which suffers from batch-to-batch variability. In contrast, purified diets are made with highly refined ingredients, and have many other advantages that make it a desirable choice [66]. Various prebiotic and probiotic supplements have been tested for modulating the gut microbiota of healthy rats, and the increase of LAB is often observed [68]. It is possible that LAB are transmitted from the gut to lungs via the gut-lung axis. Including GI samples and providing purified diets with and without supplements should be carefully designed for our future rat studies.

For ethical reasons, stage-wise multisampling in humans is mostly not feasible. Consequently, rats may serve as a valuable tool to enhance sampling and handling in microbiome studies. Unlike human beings, laboratory rats are small nocturnal caged animals, gaps and challenges exist in translating research findings gained from rat studies to human situations. Moreover, it is estimated that more than 111 million mice and rats are killed in the United States (US) laboratories each year [69]. Understanding laboratory rodents’ social life [70] and providing them with a “good life” should be prerequisites for their use [71].

## 5. Conclusions

We systematically investigated the spatial and longitudinal structures of the microbial community in 11 body compartments and across four life-history stages of healthy F344 rats. The abundances of LAB in lungs declined from breastfeed newborn to adolescence/adult and was below detectable levels in elderly rats. The lung, testes, thymus, kidney, adrenal, and muscle microbial habitats were found to have one to 52 species that changed their abundances as the rats got older. The liver and muscle exhibits the lowest microbial alpha diversity as compared to other organ systems across the four major developmental stages. Four inter-tissue microbial heterogeneity patterns were identified and characterized from integrated microbial abundance with host transcriptomic data. Breastfeeding and environmental exposure influence microbiome composition and host health and longevity. The inferred rat microbial biogeography, and the 357 pattern-specific microbial signatures, especially the LAB, should be useful for advancing human microbiome research and its translation.

## 6. List of abbreviations

SEQC: Sequencing Quality Control
LAB: Lactic acid bacteria
HMP: Human Microbiome Project
TCGA: The Cancer Genome Atlas
F344: Fischer 344
NIH: National Institutes of Health
BAP: Biology of Aging Program
GI: Gastrointestinal
FFPE: Formalin-Fixed Paraffin-Embedded
RNA-Seq: RNA sequencing
SD: Sprague Dawley
VA: Veterans Affairs
IACUC: Institutional Animal Care and Use Committee
RIN: RNA integrity number
RT-PCR: Real-time reverse transcription polymerase chain reaction
ENA: European Nucleotide Archive
TPM: Transcripts per million
BH: Benjamini-Hochberg
FDR: False discovery rate
PCoA: Principal coordinates analysis
PERMANOVA: Permutational multivariate analysis of variance
cNMF: Consensus non-negative matrix factorization
KL: Kullback-Leibler
KNN: K-nearest neighbor
KEGG: Kyoto Encyclopedia of Genes and Genomes
GO: Gene ontology
Ha-MuSV: Harvey murine sarcoma virus
Ki-MuSV: Kirsten murine sarcoma virus
OTU: Operational taxonomic unit
BMI: Body mass index
ECM: Extracellular matrix
FAO: Food and Agriculture Organization
WHO: World Health Organization
US: United States

## 7. Declarations

### 7.1 Ethics approval and consent to participate

For human data: The study was performed in accordance with the Declaration of Helsinki. The Ethics Committees of Stanford Health Care approved the study protocol. For animal data: The study had ethical approval from the Ethics Committee of IACUC.

### 7.2 Consent for publication

All three human participants signed a written informed consent.

### 7.3 Availability of data and material

The discovery dataset is available from the ENA database under accession number PRJNA238328. The newly generated rat lung data has been deposited to the ENA database under accession number PRJEB57257.

### 7.6 Authors’ contributions

Conceptualization: L.Z.; Supervision: M.R.N.; Methodology & Formal analysis: L.Z.; Animal work: C.M.C.; Sample curation: C.M.C., A.M.A.; Experiment: L.Z., C.M.C., D.K., S.G., J.L.C., M.K.A., A.M.A., K.S.; Visualization: L.Z.; Writing – original draft: L.Z.; Writing – review & editing: L.Z., M.R.N., E.S. All authors have approved the manuscript and agree with its submission.

## 9. Supplemental tables

**Table S1**. 4 microbial species abundances in 660 rat samples.

**Table S2**. 2829 species-level microbial OUT table in 660 samples.

**Table S3**. Microbial relative abundances (%) of the 4 developmental stages at the phylum, the order, and the species levels.

**Table S4**. Order-level microbial relative abundances (%) of the 11 tissues at the newborn, adolescent, adult, and senior stages.

**Table S5.** Tables of pairwise p-values of *Lactobacillales* and *Pasteurellales* from Figure 3.

**Table S6.** Age-dependent species taxonomy information and pairwise p-values between stages in six tissue types.

**Table S7.** Microbial community taxonomy information and tissue subtype classification results.

**Table S8.** Pattern-specific microbial species correlations with host genes and enriched pathways.

**Figure S1.**
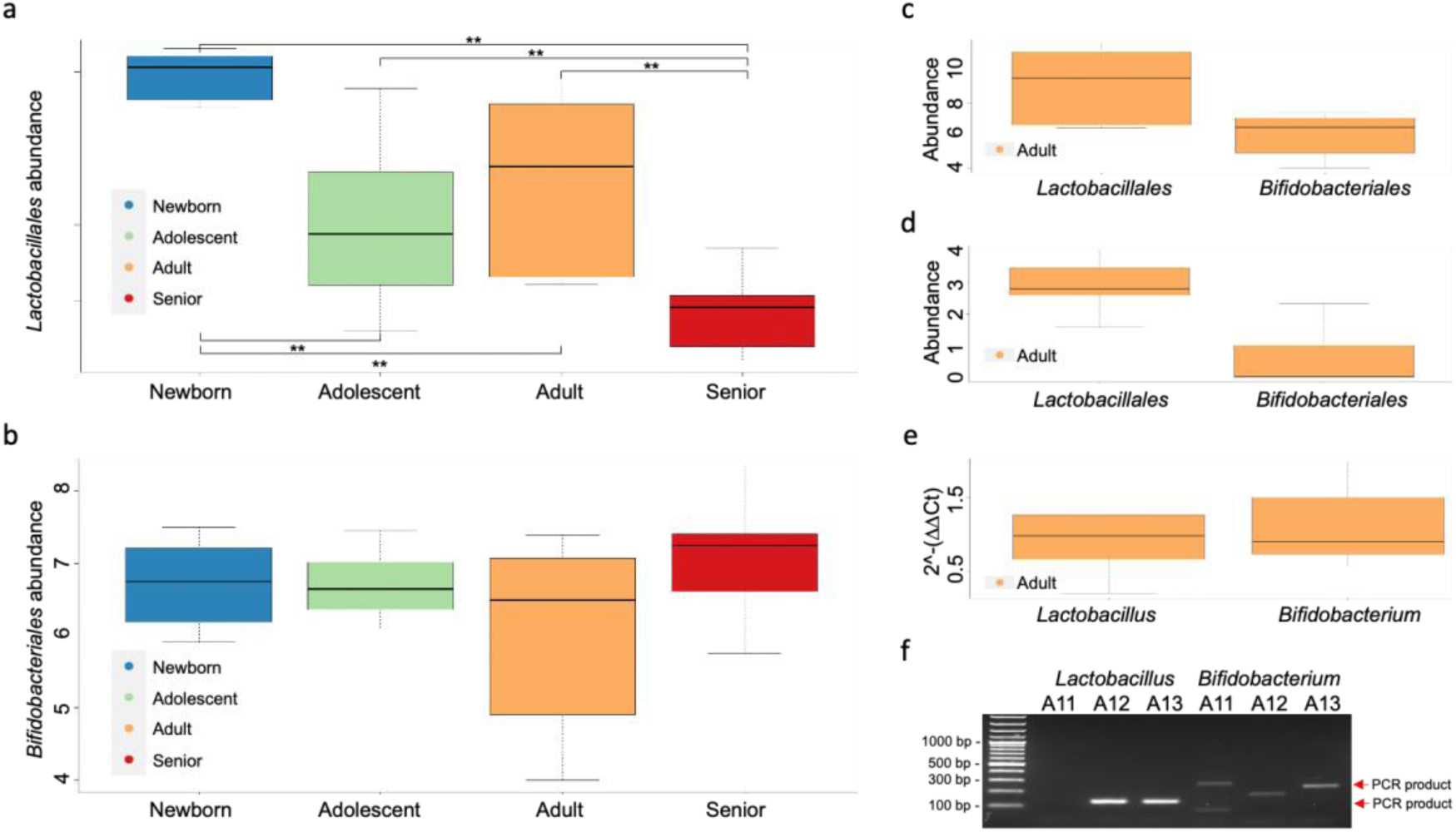
Boxplots and agarose gel showing the abundances of *Lactobacillales* and *Bifidobacteriales* in lungs at both discovery and validation datasets. (a). Boxplot illustrates the microbial abundances of *Lactobacillales* (in y-axis) at the newborn, adolescent, adult, and senior stages in the discovery dataset. (b). Boxplot illustrates the microbial abundances of *Bifidobacteriales* (in y-axis) at the newborn, adolescent, adult, and senior stages in the discovery dataset. Boxplots illustrate the microbial abundances of *Lactobacillales* and *Bifidobacteriales* (in y-axis) in the discovery (c) and validation dataset 1 (d), which were based on RNA-Seq-derived adult rats’ microbial results. Abundances in (a-d) were log2 (species counts+1). (e). Boxplot illustrates the RT-PCR microbial abundances (2^-(ΔΔCt)) in a total of 15 and 30 *Lactobacillus and Bifidobacterium* spp. (in y-axis), respectively, from the validation dataset 1. Pairwise Wilcoxon rank sum tests with BH correction were used to test for diversity differences between groups for boxplots in (a-e). Only statistically significant comparisons (p < 0.05) were marked with a single star (*), and p-values < 0.01 were marked with two stars (**). (f). Agarose gel (2%) electrophoresis of the amplified 126 bp *Lactobacillus* spp. and 100-244 bp *Bifidobacterium* spp. PCR products, respectively, in the three subjects from the validation dataset 2.

## References

1. Smith JR, Bolton ER, Dwinell MR. The Rat: A Model Used in Biomedical Research [Internet]. Methods in Molecular Biology. 2019. p. 1–41. Available from: http://dx.doi.org/10.1007/978-1-4939-9581-3_1

2. Human Microbiome Project Consortium. Structure, function and diversity of the healthy human microbiome. Nature. 2012;486:207–14.

3. Poore GD, Kopylova E, Zhu Q, Carpenter C, Fraraccio S, Wandro S, et al. Microbiome analyses of blood and tissues suggest cancer diagnostic approach. Nature. 2020;579:567–74.

4. Zhao L, Cho WCS, Luo J-L. Exploring the patient-microbiome interaction patterns for pan-cancer. Comput Struct Biotechnol J. 2022;20:3068–79.

5. Holmes DJ. F344 Rat. Sci Aging Knowledge Environ. 2003;2003:as2–as2.

6. Kwekel JC, Desai VG, Moland CL, Branham WS, Fuscoe JC. Age and sex dependent changes in liver gene expression during the life cycle of the rat. BMC Genomics. 2010;11:675.

7. Yu Y, Fuscoe JC, Zhao C, Guo C, Jia M, Qing T, et al. A rat RNA-Seq transcriptomic BodyMap across 11 organs and 4 developmental stages [Internet]. Nature Communications. 2014. Available from: http://dx.doi.org/10.1038/ncomms4230

8. Aagaard K, Ma J, Antony KM, Ganu R, Petrosino J, Versalovic J. The placenta harbors a unique microbiome. Sci Transl Med. 2014;6:237ra65.

9. Stinson LF, Boyce MC, Payne MS, Keelan JA. The Not-so-Sterile Womb: Evidence That the Human Fetus Is Exposed to Bacteria Prior to Birth. Front Microbiol. 2019;10:1124.

10. Younge NE, Araújo-Pérez F, Brandon D, Seed PC. Early-life skin microbiota in hospitalized preterm and full-term infants. Microbiome. 2018;6:98.

11. Kageyama S, Asakawa M, Takeshita T, Ihara Y, Kanno S, Hara T, et al. Transition of Bacterial Diversity and Composition in Tongue Microbiota during the First Two Years of Life. mSphere [Internet]. 2019;4. Available from: http://dx.doi.org/10.1128/mSphere.00187-19

12. Bäckhed F, Roswall J, Peng Y, Feng Q, Jia H, Kovatcheva-Datchary P, et al. Dynamics and Stabilization of the Human Gut Microbiome during the First Year of Life. Cell Host Microbe. 2015;17:852.

13. Reyman M, Clerc M, van Houten MA, Arp K, Chu MLJN, Hasrat R, et al. Microbial community networks across body sites are associated with susceptibility to respiratory infections in infants. Commun Biol. 2021;4:1233.

14. Gensollen T, Iyer SS, Kasper DL, Blumberg RS. How colonization by microbiota in early life shapes the immune system. Science. 2016;352:539–44.

15. Laursen MF, Bahl MI, Michaelsen KF, Licht TR. First Foods and Gut Microbes. Front Microbiol. 2017;8:356.

16. Faith JJ, Guruge JL, Charbonneau M, Subramanian S, Seedorf H, Goodman AL, et al. The long-term stability of the human gut microbiota. Science. 2013;341:1237439.

17. Salazar N, Valdés-Varela L, González S, Gueimonde M, de Los Reyes-Gavilán CG. Nutrition and the gut microbiome in the elderly. Gut Microbes. 2017;8:82–97.

18. Inoue R, Ushida K. Development of the intestinal microbiota in rats and its possible interactions with the evolution of the luminal IgA in the intestine. FEMS Microbiol Ecol. Oxford Academic; 2003;45:147–53.

19. Yajima M, Nakayama M, Hatano S, Yamazaki K, Aoyama Y, Yajima T, et al. Bacterial translocation in neonatal rats: the relation between intestinal flora, translocated bacteria, and influence of milk. J Pediatr Gastroenterol Nutr. 2001;33:592–601.

20. Mirpuri J, Raetz M, Sturge CR, Wilhelm CL, Benson A, Savani RC, et al. Proteobacteria-specific IgA regulates maturation of the intestinal microbiota. Gut Microbes. 2014;5:28–39.

21. Berg RD. Bacterial translocation from the gastrointestinal tract. Trends Microbiol. 1995;3:149–54.

22. Donoso F, Egerton S, Bastiaanssen TFS, Fitzgerald P, Gite S, Fouhy F, et al. Polyphenols selectively reverse early-life stress-induced behavioural, neurochemical and microbiota changes in the rat. Psychoneuroendocrinology. 2020;116:104673.

23. Luo C, Wang X, Huang H-X, Mao X-Y, Zhou H-H, Liu Z-Q. Coadministration of metformin prevents olanzapine-induced metabolic dysfunction and regulates the gut-liver axis in rats. Psychopharmacology . 2021;238:239–48.

24. Enaud R, Prevel R, Ciarlo E, Beaufils F, Wieërs G, Guery B, et al. The Gut-Lung Axis in Health and Respiratory Diseases: A Place for Inter-Organ and Inter-Kingdom Crosstalks. Front Cell Infect Microbiol. 2020;10:9.

25. Delroisse J-M, Boulvin A-L, Parmentier I, Dauphin RD, Vandenbol M, Portetelle D. Quantification of Bifidobacterium spp. and Lactobacillus spp. in rat fecal samples by real-time PCR. Microbiol Res. 2008;163:663–70.

26. Zhao L, Grimes SM, Greer SU, Kubit M, Lee H, Nadauld LD, et al. Characterization of the consensus mucosal microbiome of colorectal cancer [Internet]. Available from: http://dx.doi.org/10.1101/2021.06.02.446807

27. McMurdie PJ, Holmes S. phyloseq: an R package for reproducible interactive analysis and graphics of microbiome census data. PLoS One. 2013;8:e61217.

28. Dixon P. VEGAN, a package of R functions for community ecology. J Veg Sci. Wiley; 2003;14:927–30.

29. Adler D, Nenadic O, Zucchini W. Rgl: A r-library for 3d visualization with opengl. Proceedings of the 35th Symposium of the Interface: Computing Science and Statistics, Salt Lake City. 2003. p. 1–11.

30. Kotliar D, Veres A, Aurel Nagy M, Tabrizi S, Hodis E, Melton DA, et al. Identifying gene expression programs of cell-type identity and cellular activity with single-cell RNA-Seq [Internet]. eLife. 2019. Available from: http://dx.doi.org/10.7554/elife.43803

31. Carmona-Saez P, Pascual-Marqui RD, Tirado F, Carazo JM, Pascual-Montano A. Biclustering of gene expression data by Non-smooth Non-negative Matrix Factorization. BMC Bioinformatics. 2006;7:78.

32. Kolde R. pheatmap: Pretty heatmaps [Software]. URL https://CRAN R-project org/package=pheatmap. 2015;

33. Durinck S, Spellman PT, Birney E, Huber W. Mapping identifiers for the integration of genomic datasets with the R/Bioconductor package biomaRt. Nat Protoc. 2009;4:1184–91.

34. Kuleshov MV, Jones MR, Rouillard AD, Fernandez NF, Duan Q, Wang Z, et al. Enrichr: a comprehensive gene set enrichment analysis web server 2016 update. Nucleic Acids Res. 2016;44:W90–7.

35. Hamady M, Knight R. Microbial community profiling for human microbiome projects: Tools, techniques, and challenges. Genome Res. 2009;19:1141–52.

36. Foster T. Staphylococcus. University of Texas Medical Branch at Galveston; 1996.

37. Anderson GR, Robbins KC. Rat sequences of the Kirsten and Harvey murine sarcoma virus genomes: nature, origin, and expression in rat tumor RNA. J Virol. 1976;17:335–51.

38. Trapecar M, Leouffre T, Faure M, Jensen HE, Granum PE, Cencic A, et al. The use of a porcine intestinal cell model system for evaluating the food safety risk of Bacillus cereus probiotics and the implications for assessing enterotoxigenicity. APMIS. 2011;119:877–84.

39. Altmeyer S, Kröger S, Vahjen W, Zentek J, Scharek-Tedin L. Impact of a probiotic Bacillus cereus strain on the jejunal epithelial barrier and on the NKG2D expressing immune cells during the weaning phase of piglets. Vet Immunol Immunopathol. 2014;161:57–65.

40. Messelhäußer U, Ehling-Schulz M. Bacillus cereus—a Multifaceted Opportunistic Pathogen [Internet]. Current Clinical Microbiology Reports. 2018. p. 120–5. Available from: http://dx.doi.org/10.1007/s40588-018-0095-9

41. Isani M, Bell BA, Delaplain PT, Bowling JD, Golden JM, Elizee M, et al. Lactobacillus murinus HF12 colonizes neonatal gut and protects rats from necrotizing enterocolitis. PLoS One. 2018;13:e0196710.

42. Hu J, Deng F, Zhao B, Lin Z, Sun Q, Yang X, et al. Lactobacillus murinus alleviate intestinal ischemia/reperfusion injury through promoting the release of interleukin-10 from M2 macrophages via Toll-like receptor 2 signaling. Microbiome. 2022;10:38.

43. Yildiz S, Pereira Bonifacio Lopes JP, Bergé M, González-Ruiz V, Baud D, Kloehn J, et al. Respiratory tissue-associated commensal bacteria offer therapeutic potential against pneumococcal colonization. Elife [Internet]. 2020;9. Available from: http://dx.doi.org/10.7554/eLife.53581

44. Palmer AD, Slauch JM. Mechanisms of Salmonella pathogenesis in animal models. Hum Ecol Risk Assess. 2017;23:1877–92.

45. Eastwood J, Walton G, Van Hemert S, Williams C, Lamport D. The effect of probiotics on cognitive function across the human lifespan: A systematic review. Neurosci Biobehav Rev. 2021;128:311–27.

46. Bernig T, Taylor JG, Foster CB, Staats B, Yeager M, Chanock SJ. Sequence analysis of the mannose-binding lectin (MBL2) gene reveals a high degree of heterozygosity with evidence of selection. Genes Immun. 2004;5:461–76.

47. Kornfeld J-W, Baitzel C, Könner AC, Nicholls HT, Vogt MC, Herrmanns K, et al. Obesity-induced overexpression of miR-802 impairs glucose metabolism through silencing of Hnf1b. Nature. 2013;494:111–5.

48. Martín R, Langa S, Reviriego C, Jimínez E, Marín ML, Xaus J, et al. Human milk is a source of lactic acid bacteria for the infant gut. J Pediatr. 2003;143:754–8.

49. Fehr K, Moossavi S, Sbihi H, Boutin RCT, Bode L, Robertson B, et al. Breastmilk Feeding Practices Are Associated with the Co-Occurrence of Bacteria in Mothers’ Milk and the Infant Gut: the CHILD Cohort Study. Cell Host Microbe. 2020;28:285–97.e4.

50. Lyons KE, Ryan CA, Dempsey EM, Ross RP, Stanton C. Breast Milk, a Source of Beneficial Microbes and Associated Benefits for Infant Health. Nutrients [Internet]. 2020;12. Available from: http://dx.doi.org/10.3390/nu12041039

51. Hill C, Guarner F, Reid G, Gibson GR, Merenstein DJ, Pot B, et al. Expert consensus document. The International Scientific Association for Probiotics and Prebiotics consensus statement on the scope and appropriate use of the term probiotic. Nat Rev Gastroenterol Hepatol. 2014;11:506–14.

52. Banerjee S, Schlaeppi K, van der Heijden MGA. Keystone taxa as drivers of microbiome structure and functioning. Nat Rev Microbiol. 2018;16:567–76.

53. Zhao L, Cho WC, Nicolls MR. Colorectal Cancer-Associated Microbiome Patterns and Signatures. Front Genet [Internet]. Frontiers Media SA; 2021;12. Available from: https://www.frontiersin.org/articles/10.3389/fgene.2021.787176/full

54. Andrei A-Ş, Banciu HL, Oren A. Living with salt: metabolic and phylogenetic diversity of archaea inhabiting saline ecosystems. FEMS Microbiol Lett. 2012;330:1–9.

55. Lee H-S. Diversity of halophilic archaea in fermented foods and human intestines and their application. J Microbiol Biotechnol. 2013;23:1645–53.

56. Oxley APA, Lanfranconi MP, Würdemann D, Ott S, Schreiber S, McGenity TJ, et al. Halophilic archaea in the human intestinal mucosa. Environ Microbiol. 2010;12:2398–410.

57. Lurie-Weinberger MN, Gophna U. Archaea in and on the Human Body: Health Implications and Future Directions. PLoS Pathog. 2015;11:e1004833.

58. Zhou D, Zhang H, Bai Z, Zhang A, Bai F, Luo X, et al. Exposure to soil, house dust and decaying plants increases gut microbial diversity and decreases serum immunoglobulin E levels in BALB/c mice. Environ Microbiol. 2016;18:1326–37.

59. Blum WEH, Zechmeister-Boltenstern S, Keiblinger KM. Does Soil Contribute to the Human Gut Microbiome? Microorganisms [Internet]. 2019;7. Available from: http://dx.doi.org/10.3390/microorganisms7090287

60. Haque M, Sartelli M, McKimm J, Abu Bakar M. Health care-associated infections - an overview. Infect Drug Resist. 2018;11:2321–33.

61. Bowerman KL, Knowles SCL, Bradley JE, Baltrūnaitė L, Lynch MDJ, Jones KM, et al. Effects of laboratory domestication on the rodent gut microbiome. ISME Communications. Nature Publishing Group; 2021;1:1–14.

62. Lleal M, Sarrabayrouse G, Willamil J, Santiago A, Pozuelo M, Manichanh C. A single faecal microbiota transplantation modulates the microbiome and improves clinical manifestations in a rat model of colitis [Internet]. EBioMedicine. 2019. p. 630–41. Available from: http://dx.doi.org/10.1016/j.ebiom.2019.10.002

63. Brooks SPJ, McAllister M, Sandoz M, Kalmokoff ML. Culture-independent phylogenetic analysis of the faecal flora of the rat. Can J Microbiol. 2003;49:589– 601.

64. Manichanh C, Reeder J, Gibert P, Varela E, Llopis M, Antolin M, et al. Reshaping the gut microbiome with bacterial transplantation and antibiotic intake. Genome Res. 2010;20:1411–9.

65. Delroisse J-M, Boulvin A-L, Parmentier I, Dauphin RD, Vandenbol M, Portetelle D. Quantification of Bifidobacterium spp. and Lactobacillus spp. in rat fecal samples by real-time PCR [Internet]. Microbiological Research. 2008. p. 663–70. Available from: http://dx.doi.org/10.1016/j.micres.2006.09.004

66. Pellizzon MA, Ricci MR. Choice of Laboratory Rodent Diet May Confound Data Interpretation and Reproducibility [Internet]. Current Developments in Nutrition. 2020. Available from: http://dx.doi.org/10.1093/cdn/nzaa031

67. Pellizzon MA, Ricci MR. The common use of improper control diets in diet-induced metabolic disease research confounds data interpretation: the fiber factor. Nutr Metab . 2018;15:3.

68. Čoklo M, Maslov DR, Kraljević Pavelić S. Modulation of gut microbiota in healthy rats after exposure to nutritional supplements. Gut Microbes. 2020;12:1–28.

69. Carbone L. Estimating mouse and rat use in American laboratories by extrapolation from Animal Welfare Act-regulated species. Sci Rep. 2021;11:493.

70. Schweinfurth MK. The social life of Norway rats (Rattus norvegicus). Elife [Internet]. 2020;9. Available from: http://dx.doi.org/10.7554/eLife.54020

71. Makowska IJ, Weary DM. A Good Life for Laboratory Rodents? ILAR J. Oxford Academic; 2020;60:373–88.

